# Overcoming toxicity: why boom-and-bust cycles are good for non-antagonistic microbes

**DOI:** 10.1101/2024.08.09.607393

**Authors:** MingYi Wang, Alexander Vladimirsky, Andrea Giometto

## Abstract

Antagonistic interactions are critical determinants of microbial community stability and composition, offering host benefits such as pathogen protection and providing avenues for antimicrobial control. While the ability to eliminate competitors confers an advantage to antagonistic microbes, it often incurs a fitness cost. Consequently, many microbes only produce toxins or engage in antagonistic behavior in response to specific cues like quorum sensing molecules or environmental stress. In laboratory settings, antagonistic microbes typically dominate over sensitive ones, raising the question of why both antagonistic and non-antagonistic microbes are found in natural environments and host microbiomes. Here, using both theoretical models and experiments with killer strains of *Saccharomyces cerevisiae*, we show that boom-and-bust dynamics caused by temporal environmental fluctuations can favor non-antagonistic microbes that do not incur the growth rate cost of toxin production. Additionally, using control theory, we derive bounds on the competitive performance and identify optimal regulatory toxin-production strategies in various boom-and-bust environments where population dilutions occur either deterministically or stochastically over time. Our findings offer a new perspective on how both antagonistic and non-antagonistic microbes can thrive under varying environmental conditions.

## Introduction

Antagonistic interactions are found throughout the microbial tree of life [1–5], in almost any environment [6–10] and host-associated microbiomes [11–14]. Microbes have evolved a large variety of mechanisms to interact antagonistically with each other [1, 15], from contact dependent antagonism (e.g., via type IV, V, and VI secretion systems), to short-distance interaction mediated by diffusible antimicrobial metabolites (e.g., bacteriocins), to long-range interaction via secretion of volatile antimicrobials [16]. These antagonistic interactions are thought to be a strong determinant of microbial community structure [1, 17], to provide benefits to hosts, such as protection against pathogen invasion [13], and to be a promising avenue for antimicrobial control in both natural ecosystems and animal hosts [13, 18, 19].

Recent experimental investigations have explored the ecological [20– 22] and evolutionary dynamics of microbial antagonism [23] in the laboratory, where toxin-producing strains are found to dominate over toxin-sensitive ones. In many microbiomes, however, one finds both antagonistic and non-antagonistic microbes [24], raising the question of whether the dominance of toxin-producing strains observed in the laboratory might be caused by idealized growth conditions in those settings. Here, we explore how the interplay of environmental fluctuations, costs associated with toxin production, and regulation of toxin production, play different roles in determining the outcome of the competition. Although spatial heterogeneity of populations is one of the popular explanations of why toxin-sensitive strains might be doing well in practice [4, 25–27], we show that environmental fluctuations (e.g., dilutions) can favor toxin-sensitive strains even in spatially homogeneous populations.

Ecological theory has long recognized the significant impact of environmental fluctuations on community composition [28–31]. Changes in temperature, nutrient levels, and other abiotic factors critically shape the structure and dynamics of these communities. Such disturbances not only disrupt resident populations, possibly allowing new colonizers to invade, but also affect both immediate and long-term ecological outcomes [32, 33]. Microbial communities in a diverse array of habitats are subject to “boom-and-bust” dynamics, where periods of rapid population growth are often followed by sharp declines. Such dynamics have been observed across various environments, including phytoplankton and particle-attached microbial communities in marine ecosystems [34–36], soil [37–40], host-associated microbiomes [41–44], and the built environment [45]. These boom-and-bust cycles are not only caused by abiotic factors such as nutrient availability and environmental disturbances, but also by biotic interactions including competition, predation, and parasitism. Particularly, the interactions with phages and predators can drastically alter microbial community structure, trigger population crashes, and thereby influence the overall dynamics of microbial ecosystems [46– 48].

Despite the interest in the effect of microbial antagonism and environmental fluctuations on microbial community composition [33, 49, 50], the interplay between environmental fluctuations and antagonistic microbial interactions is surprisingly underexplored. Although data relating the relative abundance of toxin-producing strains to environmental fluctuations is very scarce, there is some evidence that environments with higher turnover rates harbor a reduced number of toxin-producing strains [51, 52]. Mathematical modeling and laboratory experiments in stationary environments suggest that the efficacy of toxin-mediated killing is dependent on high population densities [21, 53] and consequently frequent population busts could favor sensitive strains that avoid the metabolic costs associated with toxin production. Certain microbial species have evolved regulatory mechanisms for toxin production that are triggered by quorum sensing signals [53, 54] or environmental indicators of stationary phase [55]. This regulation ensures that resources are not expended on toxin production at times when it would be least effective, possibly allowing microbes to optimize the cost-benefit ratio of toxin production. Activation of toxin production genes in response to quorum sensing signals produced by other strains, a phenomenon known as eavesdropping or cross-talk, has also been reported [56]. Regulation of toxin production in response to self and non-self abundances is thus theoretically possible and may be exploited to design synthetic genetic systems for pathogen eradication via targeted secretion of antimicrobials [57].

Our paper seeks to investigate how boom-and-bust cycles and toxin regulation affect the antagonistic competition dynamics between toxin-producing and non-producing microbes. This approach aims to elucidate the survival strategies of microbial populations and provide a deeper understanding of the ecological and evolutionary consequences of microbial interactions under variable environmental conditions.

### Experimental antagonism with periodic dilutions

To gain intuition for the impact of environmental disturbances on the dynamics of microbial antagonism, we conducted competition experiments between a sensitive (S) and a killer (K) strain of *Saccharomyces cerevisiae*, with the latter engineered to secrete the killer toxin K1 expressed from the galactose-inducible promoter *P*_*GAL*1_ [21]. The toxin K1 kills S cells by disrupting ion balance and membrane potential [58], but does not affect K cells that are immune to it [59].

In isolation and at low density, strain S grew at a faster growth rate (*r*_*S*_ = 0.28 ± 0.01 h^−1^, mean ± SD) than strain K (*r*_*S*_ = 0.26 ± 0.03 h^−1^). Making these two strains compete against each other in well-mixed liquid cultures, we found that, for the lowest initial population sizes tested, the fraction of K cells in the population initially decreased and then increased (the lightest curves in Fig. 1). Increasing the initial population size or the initial fraction of K cells reduced the time window over which their fraction decreased. These dynamics are consistent with the intuition that at low cell densities, when the toxin is too dilute to significantly impact S cells, their relative abundance should temporarily increase due to their higher growth rate.

**Figure 1.**
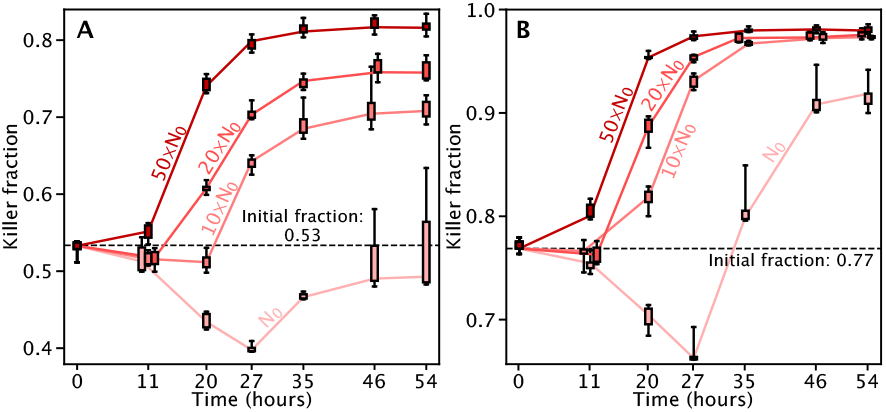
Uninterrupted competitions between the killer and sensitive strains in experiments initialized with different population sizes (different shades, *N*_0_ = 5.8 · 10^4^ cells/mL) and killer fractions *f*_0_ (53% in A and 77% in B). With small initial populations, the fraction of killers first decreases, due to the sensitive strain’s higher growth rate, and later increases, due to the toxin’s action. Lines connect subsequent median fractions. Some data points where spaced horizontally to ease visualization. Quartiles were computed via bootstrapping, whiskers report the minimum and maximum values. Could the toxin-sensitive microbes become dominant in the long term if recurrent environmental disturbances dilute the population when they are in the majority?

These observations led us to hypothesize that frequent and fortuitously timed disturbance events, such as dilutions, could favor the sensitive strain allowing it to become dominant over time, despite the presence of the toxin-producing killer strain, which would normally dominate without such dilutions. To test this hypothesis, we made the two strains compete against each other by periodically diluting them with different dilution intervals (Fig. 2). Unlike in the experiment of Fig. 1, here nutrients were replenished at each dilution, allowing the population to re-grow (Fig. S1). In agreement with our hypothesis, we found that both the initial fraction and the interval between successive dilutions influenced the competition outcome between S and K, favoring S when dilutions occurred more frequently (Fig. 2). While some heterogeneity of competition outcomes was observed with the 48 h cycle, the overall trend was clear: when the initial K fraction was low enough, S ultimately dominated, with the frequency of dilutions influencing only the rate at which S became dominant; when the initial K fraction was much higher, the same was true about the K domination; but for intermediate initial K fractions, the frequency of dilutions became a crucial predictor of the ultimate winner.

**Figure 2.**
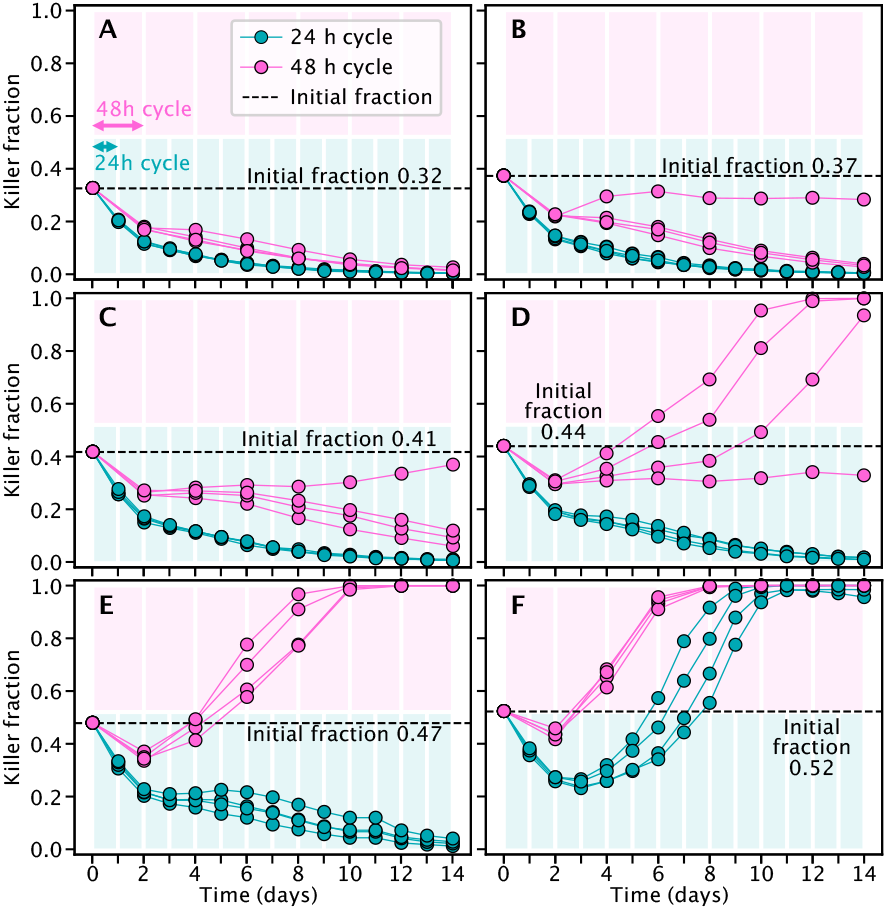
Experimental competitions between a toxin-producing (killer) and a sensitive strain of *Saccharomyces cerevisiae* in environments diluted periodically with periods of *T* = 1 day (teal) and *T* = 2 days (pink). Different panels show different initial fractions of the killer strain. Both the initial killer fraction and the period of the dilution cycles determine the outcome of the competition and the rate of extinction of the losing strain, with longer inter-dilution times favoring the killer strain. Shaded pink and teal rectangles display uninterrupted periods of growth in the two treatments. White vertical lines depict dilution events.

Overall, these experiments suggest that frequent dilutions can favor sensitive cells when their growth rates in isolation are larger than those of the killers. However, several factors may limit the general applicability of these results. For instance, many antagonistic mi-crobes regulate toxin production through quorum sensing or in response to environmental cues, rather than producing toxins constitutively. Additionally, nutrient turnover in more realistic environments could enable toxin-producers to completely eliminate toxin-sensitive microbes between population busts, unlike in our batch culture experiments where the relative proportions of K and S stabilize once nutrients are depleted (Fig. 1). To more comprehensively characterize how growth rates, toxin regulation, and environmental fluctuations jointly affect the dynamics of microbial antagonism, we turn to mathematical modeling.

### Population dynamics

We start by describing a basic competition model between a toxin-producing killer (K) and a toxin-sensitive strain (S). The goal is to keep the modeled mechanisms general, although our main ideas can be similarly applied to more complex models tied to specific microorganisms, experimental conditions, or environments.

We assume that the growth of both strains is logistic, with respective intrinsic growth rates (*r*_K_, *r*_S_) and a shared carrying capacity^1^ *C*. We assume that the killer’s *ability* to produce the toxin confers to it a growth rate deficit (i.e., *r*_K_ < *r*_S_) independent of the actual toxin production. This deficit could arise from costs related to the regulation of toxin production, such as quorum sensing [60], but may also reflect other metabolic or genotypic differences that are unrelated to toxin production. Additionally, we assume an extra growth rate deficit that scales linearly with the normalized toxin production rate *a* ∈ [0, 1], yielding the realized growth rate *r*_K_(1 − *εa*), where *ε* > 0 is the cost of producing the toxin at the maximal rate. This deficit accounts for the metabolic cost of toxin production [53] and other expenses, such as toxin secretion, which may require cell lysis [3]. The rate of toxin-induced death of the sensitive strain is assumed to be proportional to the product of the strain densities (*n*_K_ and *n*_S_, respectively) with the killing rate *µ*. After non-dimensionalizing, *n*_K_(*t*) → *n*_K_(*t*)/*C, n*_S_(*t*) → *n*_S_(*t*)/*C*, and *t* → *r*_S_*t*, the resulting dynamics are

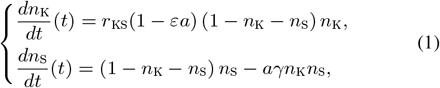

where *r*_KS_ = *r*_K_/*r*_S_ is the intrinsic growth rates ratio and *γ* = *µC/r*_S_ is the rescaled killing rate. In our numerical experiments, we set *γ* = 1 so that the strength of antagonism (the toxin-induced death term − *γn*_K_*n*_S_ with the maximum rate of toxin-production *a* = 1) is comparable to the inter-strain competition for nutrients (the quadratic term −*n*_K_*n*_S_ in the logistic). For the sake of consistency, all rates in the rest of this paper are dimensionless (i.e., scaled by *r*_S_).

Distinguishing between the relative values of *r*_KS_ and *ε* in published datasets is challenging, as both parameters influence the realized growth rate of the killer strain. A review of various studies [61– 63] that compared the growth rates of killer and sensitive strains suggests that the ratio of realized growth rates *r*_KS_(1 −*ε*) typically falls between 0.68 and 0.98 for strains that produce the toxin at a constant, maximal rate (referred to as *constitutive killers*). Since proteinaceous toxins are often expressed from plasmids, one can estimate characteristic *r*_KS_ values by comparing the growth rates of *Escherichia coli* strains harboring such plasmids, with the toxin-producing genes deleted, to strains lacking these plasmids. From [64], we derive that *r*_KS_ ≈ 0.85 is a plausible value, based on comparisons of growth rates between cells with and without ColE1-type plasmids. In our experiments with killer *S. cerevisiae*, the cost of constitutive toxin production (*ε*) is minor [21], but the deletion of the hexokinase isoenzyme 2 gene that allows expression of the toxin from the galactose-inducible promoter *P*_*GAL*1_ in K carries a growth-rate cost that reduces *r*_K_ compared to *r*_S_, resulting in the ratio of realized growth rates *r*_K_(1 − *ε*)/*r*_S_ = 0.92 ± 0.03 (mean±SE). Unless otherwise noted, we will adopt *r*_KS_ = 0.85 and *ε* = 0.2 in our computations, aligning with the lower bound *r*_KS_(1 − ε) = 0.68 of the range reported in the literature.

#### Box 1: Population growth model

Switching from the normalized strain abundances (*n*_K_, *n*_S_) to the normalized population size (*N* (*t*) = *n*_K_(*t*) + *n*_S_(*t*) ∈ [0, 1]) and the fraction of killers (*f* (*t*) = *n*_K_(*t*)/*N* (*t*) ∈ [0, 1]), we re-write Eq. (1) as

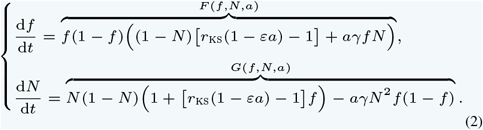

Under this transformation, the entire horizontal line *N* = 0 maps to the origin (*n*_K_, *n*_S_) = (0, 0) in the original Eq. (1) coordinates. Similarly, the horizontal line *N* = 1 corresponds to *n*_K_ + *n*_S_ = 1, indicating that the system is at carrying capacity.

**Definitions and Parameters:**

- *f* (0) = *f*_0_, *N* (0) = *N*_0_: initial killer fraction and population size;
- *r*_KS_ := *r*_K_/*r*_S_: ratio between intrinsic growth rates;
- *a*(*t*) ∈ [0, 1]: toxin-production rate;
- *ε* > 0: cost of producing the toxin;
- *γ* = *µC/r*_S_: rescaled killing rate.

It is also more convenient to restate the dynamics in terms of the normalized total population size *N* (*t*) = *n*_K_(*t*) + *n*_S_(*t*) and the fraction of killers *f* (*t*) = *n*_K_(*t*)/*N* (*t*). This change of coordinates yields the ODE model Eq. (2) on a unit square, as summarized in Box 1. Fig 3A shows the phase portrait for constitutive killers, with all trajectories approaching *f* = 1 and *N* = 1 (i.e., (*n*_K_, *n*_S_) = (1, 0) – the competitive exclusion of sensitives by killers), which is the only attracting fixed point of Eq. (2) for any fixed *a* > 0. Consistently with experimental data in Fig. 1, when starting from a small initial population *N*, the trajectories in Fig 3A bend left (i.e., decreasing the killer fraction) for a significant amount of time. This reduction is due to the killers’ growth-rate disadvantage (*r*_KS_(1 − *ε*) < 1), which does not prevent their eventual domination.

**Figure 3.**
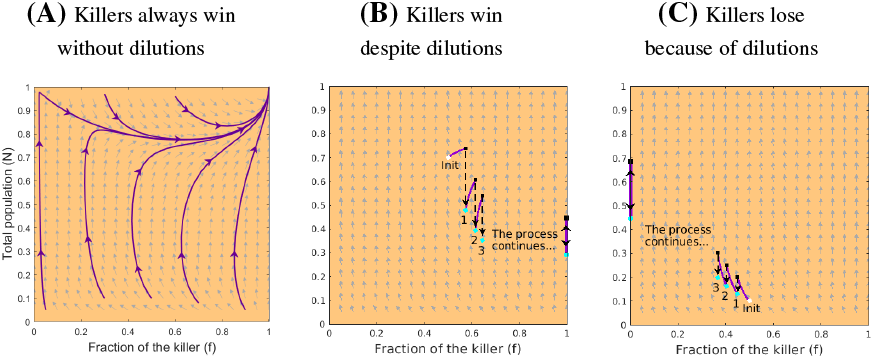
Trajectories of competitions between constitutive killers and sensitives in undisturbed (A) and periodically-diluted (B,C) populations. (A) Killers always win without dilutions. (B,C) With periodic dilutions, their fate depends on the initial condition. (B) With a high enough initial population size, e.g., (*f*_0_, *N*_0_) = (0.5, 0.7), a sequence of dilutions carries killers to an eventual victory (i.e., *f* → 1). (C) Starting at a lower population size, e.g., (*f*_0_, *N*_0_) = (0.5, 0.1), the dilutions lead to their demise (i.e., *f* → 0). In both cases, the period of dilutions is *T* = 1, and the pre- and post-dilution states are shown with black squares and cyan dots respectively. Once either strain dominates, the population oscillates between the terminal cyan dot and black square at *f* = 0 or *f* = 1. Temporal trajectories associated with panel B are shown in Fig. 4B. Gray arrows denote the vector field directions corresponding to Eq. (2) with *a* = 1. Parameter values: *ε* = 0.2, *r*_KS_ = 0.85, *γ* = 1, and *ρ* = 0.65 used here and throughout the paper unless stated otherwise.

### Do regular dilutions protect the sensitive?

We begin by assuming that dilutions occur regularly every *T* time units and that at each dilution the relative strain abundances are preserved, but only a fixed fraction *ρ* of the total population survives^2^. We observe that with relatively frequent dilutions (*T* = 1, i.e., the inter-dilution time equal to the inverse growth rate of sensitive cells), the killers may either progressively increase their relative abundance (Fig. 3B) or decrease it (Fig. 3C), depending on the initial condition.

This suggests that while dilution events can disrupt the killers’ dominance, a further investigation is needed to determine under which circumstances these interventions help the sensitives if *T* is small.

Numerical methods make it easy to analyze the performance of constitutive killers for all possible initial states over one cycle; i.e., we use linear partial differential equations (PDEs) to compute the pre-dilution *f* (*T* ^−^) = *f* (*T* ^+^) and *N* (*T* ^−^) = *N* (*T* ^+^)/*ρ* corresponding to all initial (*f*_0_, *N*_0_); see Fig. 4A and SI Appendix §S7.1. Once this mapping is computed, we iterate it to determine the asymptotic outcomes as the number of dilutions approaches infinity. Fig. 4A shows a general trend: larger initial killer fractions *f*_0_ and population sizes *N*_0_ allow constitutive killers to moderately increase their relative abundance by the end of the first cycle. The region below the black-dashed curve in Fig. 4A indicates initial conditions for which killers decrease their fraction in the first cycle; i.e., *f* (*T* ^−^) < *f*_0_. Thus, one may expect that populations that start below the black-dashed curve are exactly the ones that become dominated by the sensitive strain with successive dilutions. However, our calculation of the asymptotic limit 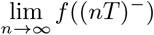, shows that this is not the case. Fig. 4C-D show that, in the limit of infinite dilutions, the state space is divided into two regions corresponding to the competitive exclusion of the killer by the sensitive strain (blue) and vice versa (red). The actual shades of blue and red in Figs. 4C-D represent the time it takes from the initial (*f*_0_, *N*_0_) to come within the machine accuracy of the asymptotic limit for killers’ pre-dilution fraction (which in real systems would be correlated with the time until the competitive exclusion). We highlight five features generic in these computations, which mirror the observations from the experiments of Fig. 2:

**Figure 4.**
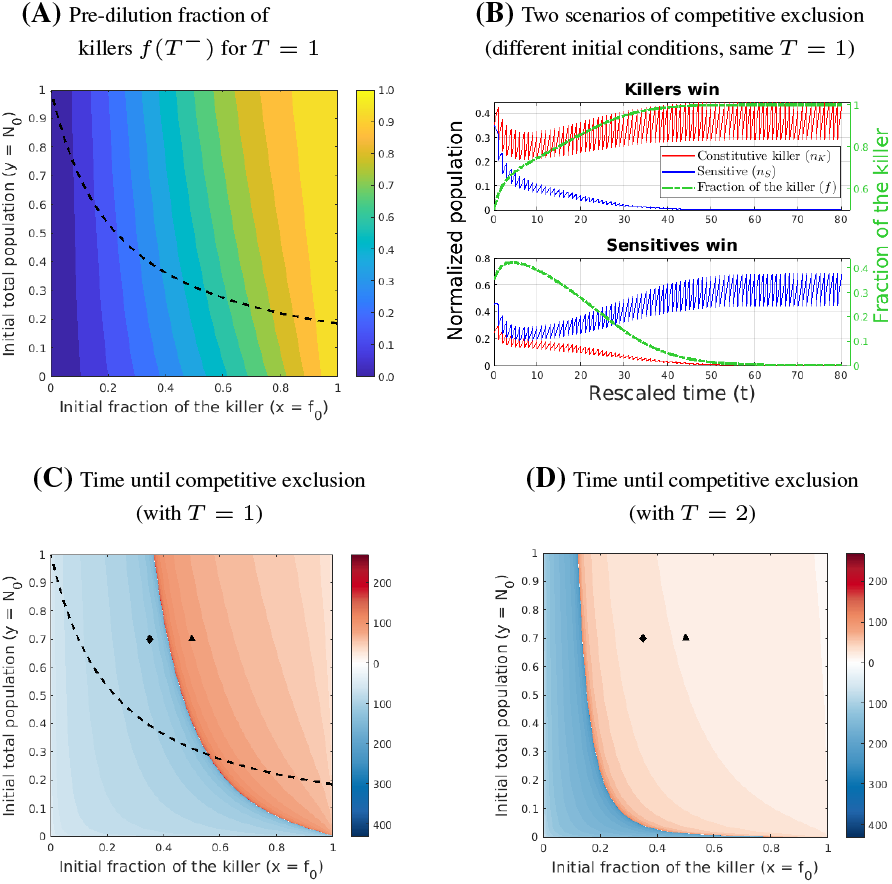
Constitutive killers with regular dilutions: pre-dilution fraction and limiting behavior. (A) Pre-dilution fraction of the killer at the end of the first cycle, *f* (*T* ^−^), for *T* = 1 and any initial condition. Initial conditions below the black, dashed curve lead to *f* (*T* ^−^) < *f*_0_. (B) Temporal trajectories of killer (red) and sensitive (blue) population sizes (left axes), and killer fraction (green, right axes), with two different initial conditions (corresponding to the ♦ and ▴ markers in panel C) and dilution period *T* = 1. (C) Time until competitive exclusion (red/blue shades) and limiting killer fraction (red and blue indicate *f* = 1 and *f* = 0 limits, respectively) vary with the initial condition. Within the red and blue regions, the absolute killer and sensitive population sizes reach *n*_K_ ≈ 0.45 and *n*_K_ ≈ 0.69, respectively (see also panel B). Black dashed line as in panel A. (D) Doubling the dilution period extends the range of initial conditions leading to domination by the killer and reduces/increases the timescale over which the killer/sensitive reach domination, respectively. Both initial conditions marked by ♦ and ▴ now lead to killer domination.

1. For every initial population size, there exists a critical killer fraction threshold that determines which strain will eventually dominate.
2. The closer the initial killer fraction is to that threshold, the longer it takes to approach competitive exclusion.
3. Changes in relative abundance in the first cycle are not predictive of the asymptotic limit. Accordingly, in our experiments of Fig. 2, the killer’s fraction always decreased in the first cycle due to the low initial population size.
4. When the dilution period grows, this might change the asymptotic limit and the time necessary to approach it. For example, the ♦-marked initial condition leads to the victory of sensitives when *T* = 1 and of killers when *T* = 2 (Fig. 4C-D). While the ▴-marked initial condition was already leading to killers’ victory even with *T* = 1, with *T* = 2 this exclusion happens much faster. This is consistent with the comparison of 24 h and 48 h dilution trajectories in the experiments.
5. With *T* = 1, neither strain can reach the carrying capacity within one cycle even after the other strain is excluded in the limit. The range of such oscillations can be found analytically (SI Appendix §S3.1), is observed in Fig. 4B, and also consistent with experimental evidence in Fig. S1.

### Do toxin-producers benefit from population-sensing?

The results for constitutive killers are revealing and align well with our experiments. However, antagonistic strains often use quorum sensing or environmental signals [53–55] to regulate toxin production, and thus may not engage in antagonistic behavior at all times, decreasing their growth-rate penalty when the toxin is not produced. Here, rather than modeling specific types of quorum sensing mechanisms, we use a phenomenological approach and explore different notions of optimality for toxin production policies of “omniscient” killers. The results will serve as an upper bound on how well more realistic killers could do given the limits to their sensing abilities.

Supposing that the killers can sense the current size and composition of the population, they could use this to regulate their rate of toxin production, also taking into account the remaining time until the next dilution. More precisely, we will consider a theoretical possibility of their evolving the optimal toxin production rate *in feedback form*: *a*_*_ = *a*_*_(*f, N, t*) to optimize the resulting pre-dilution fraction *f* (*T* ^−^). We will refer to such killers as “myopically-optimal” or simply “myopic” since they optimize the results over a single cycle only, without any regard to the sequence of future dilutions. We use the methods of control theory [65] to find this optimal toxin-production policy in the framework of dynamic programming;^3^ see the summary in Box 2 and computational details in SI Appendix §S7. The structure of this control problem guarantees that the optimal policy is *bang-bang*; i.e., for generic (*f, N, t*), it will be optimal to either not produce the toxin at all (*a*_*_ = 0) or produce it at the maximum rate (*a*_*_ = 1).

#### Box 2: Myopic optimal toxin production policy for population-sensing killers under regular dilutions

The *value function* 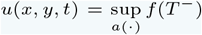 is the best pre-dilution killer fraction achievable starting from *f* (*t*) = *x, N* (*t*) = *y*. This value function satisfies a Hamilton-Jacobi-Bellman (HJB) PDE

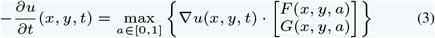

with terminal conditions *u*(*x, y, T*) = *x* if *y* > 0 and *u*(*x*, 0, *T*) = 0. *F* and *G* are defined in Eq. (2). The myopically optimal toxin-production policy *a*_*_(*x, y, t*) is an argmax in Eq. (3), see SI Appendix §S7.1.

The corresponding pre-dilution population size *ϕ*(*x, y, t*) = *N* (*T*^−^) starting from *f* (*t*) = *x, N* (*t*) = *y*, and using policy *a*_*_(·) satisfies

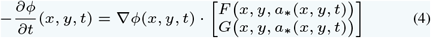

with the terminal condition *ϕ*(*x, y, T*) = *y*.

Iterating (*u, ϕ*) one can obtain the limiting pre-dilution fraction of killers 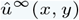 under an infinite sequence of dilutions (SI Appendix §S7.2).

As Fig. 5C shows, these myopic killers try to maximize the early exponential growth by opting not to produce the toxin at first if the initial population and/or their initial fraction are low. Fig. 5A demonstrates that they do better than the constitutive killers over the first cycle, but at least for these parameter values, this sensing-based advantage is minor and mostly pronounced when the initial populations are low. Compared to constitutive killers, the region starting from which the myopic killers eventually dominate expands only slightly, for small *N*_0_ and relatively large *f*_0_ (Fig. 5B). Consequently, the sensitive strain is still protected by periodic dilutions, which allow it to achieve competitive exclusion starting from a broad range of initial configurations.

**Figure 5.**
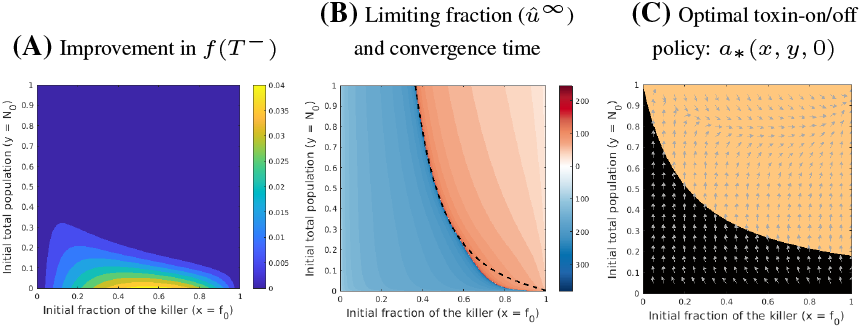
Myopic killers with regular dilutions: pre-dilution and limiting behaviors with regular dilutions and *T* = 1. (A) Population sensing and myopic planning provide a minor improvement to the killer fraction *f* (*T* ^−^) at the end of the first cycle, mostly for initial conditions with low population size. Shown here is the maximized *f* (*T* ^−^) for myopic killers, minus the corresponding *f* (*T* ^−^) for constitutive killers. (B) In the infinite-dilution limit, myopic population sensing expands the set of initial conditions leading to the killer dominance, compared to constitutive killers (black dashed curve reports the blue/red boundary of Fig. 4C). (C) The optimal toxin production policy *a*_*_(*f, N, t*) (shown here for the initial time *t* = 0 only) is *bang-bang*, equal to 1 in the orange region, and to 0 in the black region. Gray arrows denote the vector field directions corresponding to Eq. (2) with *a* = *a*_*_(*f, N*, 0). When the killers switch from no toxin production to constitutive production (i.e., as the competition trajectory crosses from the black to orange region in (*f, N*) space), the vector field exhibits a jump discontinuity. However, the angular difference between the vectors on either side is consistently small (always < 1.27°), making the discontinuity not visually obvious. The policy is time-dependent and the toxin production region changes for *t* > 0.

### Who benefits from randomness in dilution times?

The periodic setting investigated up till now provides useful insights but may oversimplify the complex dynamics of populations in natural environments. For example, boom-and-bust dynamics are often driven by fluctuations that occur randomly in time, rather than periodically. Here, we extend our analysis to randomly distributed perturbations.

We model dilutions as random events governed by a Poisson process; so, the duration of inter-dilution time intervals T are exponentially distributed with rate *λ* > 0. As before, we will consider proportional dilutions, preserving *f* but instantaneously switching from *N* to *ρN*. Mathematically, this continuous evolution of *f* (*t*) and *N* (*t*) punctuated by randomly timed jumps in *N* can be described as a Piecewise-Deterministic Markov Process (PDMP) [67] and we take advantage of a well-developed theory for optimal control of PDMPs throughout the rest of this paper. We first note that any toxin-production policies will now be independent of time since the last dilution; we will use *α* = *α*(*f, N*) to denote such feedback policies, to distinguish them from *a*(*f, N, t*) used in the periodic case above.

Second, we note that our criteria for evaluating initial conditions and the quality of policies become more subtle. Starting from the same initial (*f*_0_, *N*_0_) and using any reasonable fixed policy *α*, a sequence of randomly timed dilutions might lead to a competitive exclusion of either strain. Fig. 6 illustrates this for the simplest case of constitutive killers; from now on, we will use *α*_0_ = 1 to denote their policy. Since a trajectory reaches neither *f* = 0 nor *f* = 1 in finite time, we will select small threshold values *γ*_d_ and (1 − *γ*_v_), declaring killers’ victory^4^ as soon as *f* (*t*) > *γ*_v_ or sensitives’ victory as soon as *f* (*t*) < *γ*_d_. We then define a new probabilistic metric for policy performance: 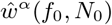 is the probability of the killers attaining their victory (before the toxin-sensitives do) after an arbitrary number of dilutions starting from (*f*_0_, *N*_0_) and using policy *α*. The rigorous definition and the PDE that 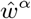 must satisfy are covered in Box 3. We will use this metric to compare the performance of all toxin-production policies in stochastic environments.

**Figure 6.**
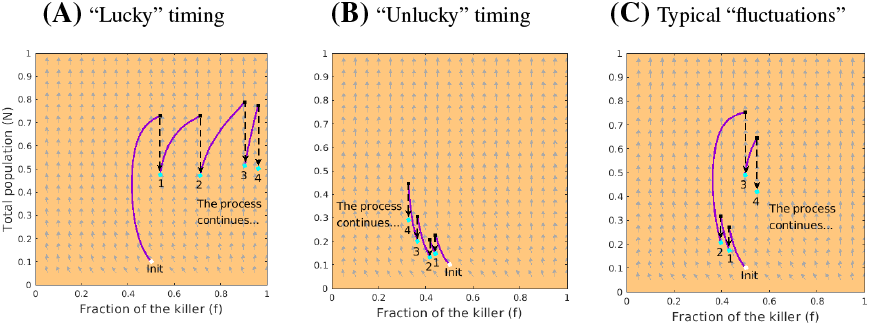
Constitutive killers with random dilution times. Killers can either progressively increase or decrease their fraction in a “lucky” or “unlucky” scenario depicted in panels (A, B), respectively. However, in most cases, their fraction will fluctuate instead (panel C). In all cases, the initial configuration (*f*_0_, *N*_0_) = (0.5, 0.1) is plotted with a white dot and followed by 4 dilution events. The pre-dilution / post-dilution states are again shown with black squares and cyan dots, respectively. Gray arrows denote the vector field directions corresponding to Eq. (2) with *a* = 1.

#### Box 3: Probabilistic performance metric for policies under randomly-timed dilutions

**Definitions and Parameters:**

- Δ_v_ = {(*x, y*) ∈ [0, 1]_2_ | *x* > *γ*_v_ }, victory zone (*γ*_v_ –victory threshold);
- Δ_d_ = {(*x, y*) ∈ [0, 1]_2_ | *x* < *γ*_d_ }, defeat zone (*γ*_d_ –defeat threshold);
- Δ = Δ_v_ ∪ Δ_d_, terminal set.

**(Random) victory time for the killer:**

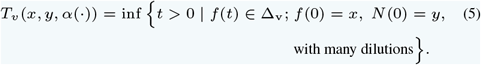

with many dilutions}.

**(Random) defeat time for the killer:**

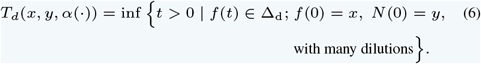

**(Random) termination time:**

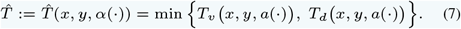

**Terminal cost:**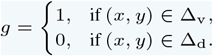,

**Performance metric (probability of killers’ victory with** *α*):

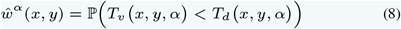

can be found by numerically solving a first-order linear equation:

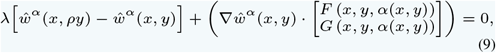

with the boundary condition 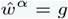 on Δ.

See **Remark VII** in SI Appendix §S7.3 for the numerics.

Since the inter-dilution intervals are random, this also affects the notion of optimal policy. Extending the same myopic approach to stochastic setting, we will call the killers *tactically-optimal* if they follow a policy *α*_1_ chosen to maximize the *expected* killers’ fraction just before the next dilution; i.e., 𝔼 [*f* (𝒯^−^)]. The subscript in *α*_1_ indicates the temporal horizon for optimizing this policy (one cycle). This bang-bang policy can be found by solving the so-called “randomly-terminated” problem [68]; mathematical details are in Box 4, part 2.

#### Box 4: Different types of toxin production policies under randomly-timed dilutions

1. **Constitutive policy** *α*_0_ : always produce the toxin; i.e., *α*_0_ (*f, N*) = 1.
2. **Tactically-optimal policy** *α*_1_ : Maximizes the *expected* killers’ fraction just before the next dilution. Given the value function

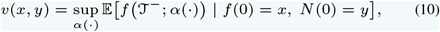

the tactically-optimal policy *α*_1_ can be found by solving a first-order HJB PDE satisfied by *v*:

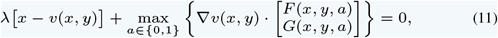

with the boundary condition

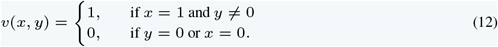 See SI Appendix §S6.1 for the derivation.
3. **Strategically-optimal policy** *α*_∞_ :

Maximizes the probability of killers winning before the sensitives do (without any limit on the number of dilutions). Given the value function

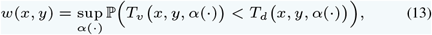

the strategically-optimal policy *α*_∞_ can be found by solving a first-order non-local HJB equation satisfied by *w*:

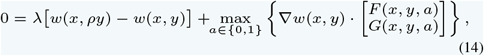

with the boundary condition

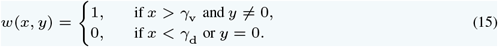

See SI Appendix §S6.2 for the derivation and §S7.3 for the numerics.

We first focus on the case *λ* = 1, to ensure that 𝔼[𝒯] = 1/*λ* = 1 matches the period of regular dilutions *T* = 1 considered in the previous section. Fig. 7 compares the performance of constitutive and tactically-optimal killers. Unlike in the periodic case (Fig. 5A), here the advantage of population-sensing (tactically-optimal) killers is significant: they have noticeably better chances of winning than constitutives starting from most initial conditions. Another simple comparison is to focus on the previous boundaries between the blue (deterministic defeat) and red (deterministic winning) in Figs. 4C and 5B. Plotting these boundaries as black dashed lines in Figs. 7A and 7B respectively, we provide a different quantitative measure of tactically-optimal killers’ advantage: their average chances for success starting near this deterministic “no microbe’s land” are ≈ 72%, compared to only ≈ 46% for the constitutive killers. Interestingly, the tactically-optimal policy *α*_1_ (*f, N*) shown in Fig. 7C is quite close to the zeroth time-slice of the deterministic myopic optimal policy *a*_*_(*f, N*, 0) from Fig. 5C. The difference in performance comes primarily from the fact that *α*_1_ (*f, N*) is stationary and that occasional long intervals (𝒯 *>* 1/*λ*) really help the tactically-optimal killers. Nevertheless, the toxin-sensitives still have a significant probability of winning on a large set of initial conditions, particularly when the toxin-producers are not starting in the majority.

**Figure 7.**
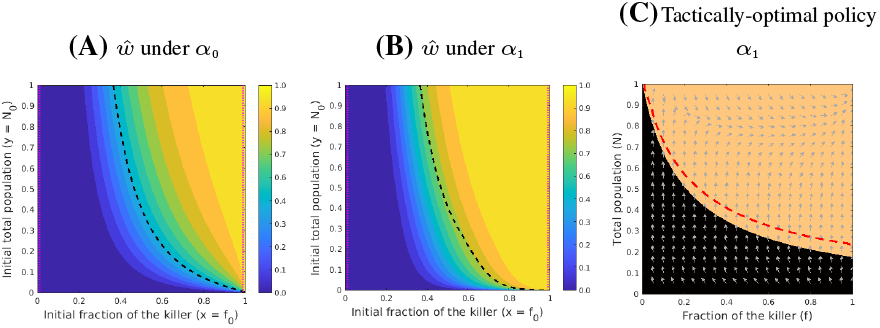
Probabilistic performance of (A) constitutive and (B) tactically-optimal killers with random dilution times (*λ* = 1). The probability of attaining competitive exclusion is noticeably higher for the tactically-optimal toxin-producers starting from most initial configurations. Dashed black lines show the boundary of the set from which they could (deterministically) win under periodic dilutions with *T* = 1. In the current random dilutions setting, starting near that boundary gives the tactically-optimal killers an average ≈ 72% chance of winning, compared to only ≈ 46% for the constitutive killers. This is mainly because the tactically-optimal killers do not produce the toxin when their fraction or the overall population size is low (black region in C). In both (A) & (B), the victory and defeat barriers (*γ*_v_ and *γ*_d_, respectively) are indicated by vertical magenta dotted lines. In (C), gray arrows represent the vector field directions corresponding to Eq. (2) with *a* = *α*_1_ (*f, N*), and the red-dashed line denotes the lower boundary of the toxin-on region for a different *strategically-optimal* policy *α*_∞_ discussed below. All parameter values are the same as in Fig. 4C.

For each toxin production policy, it is also interesting to ask whether it performs better in deterministic or stochastic environments (assuming that the average dilution frequency and strength are the same in both cases). In the SI Appendix §S4, we perform this comparison by fixing a specific initial population size *N*_0_ = 0.5 and averaging over possible initial killer fractions *f*_0_. The observed perfor-mance differences depend non-trivially on *ρ* and *λ*; see Fig. S6. But in general, constitutive killers perform better in stochastic environments when *λ* is large and *ρ* is small, and in deterministic environments when *λ* is small and *ρ* is large. In contrast, tactically-optimal population-sensing killers always fare better under stochastic dilutions.

### Can toxin-producers do better if they are non-myopic?

The optimality of toxin-production policy *α*_1_ is tactical (or myopic) because it is selected with only one upcoming dilution in mind, ignoring the ultimate goal of killers to win after arbitrarily many dilutions. It is natural to ask whether they would gain a substantial advantage by selecting a policy that maximizes the probability of attaining their victory before the sensitives, 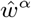. For fixed values of dilution frequency *λ* and survival factor *ρ*, such *strategically-optimal* policy *α*_∞_ (*f, N*) can be computed as described in Box 4 part 3. The subscript in *α*_∞_ indicates that the policy maximizes the probability of eventual victory without any regard to how many dilutions will be required. The strategically-optimal *α*_∞_ prescribes producing toxin slightly more conservatively than the tactical *α*_1_ (see the red-dashed line in Fig. 7C). But in the end, *α*_∞_’s performance is only marginally better for our chosen parameter values; see SI Appendix §S5 and also Fig. 8C.

**Figure 8.**
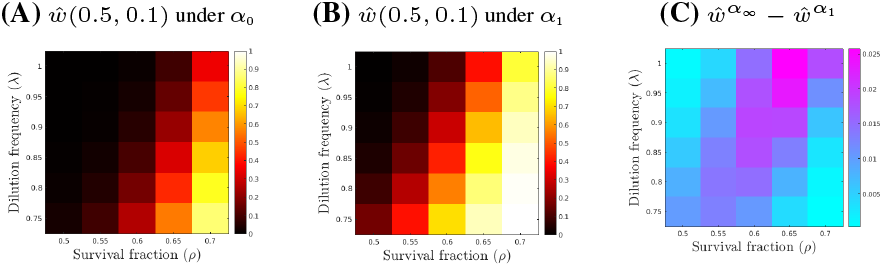
Comparison of probabilistic performance for different toxin production policies starting from (*f*_0_, *N*_0_) = (0.5, 0.1) for a range of survival fractions and dilution frequencies, with random dilution times. Policy *α*_1_ is recomputed for each *λ*, while policy *α*_∞_ is recomputed for each (*ρ, λ*) combination. In general, a larger survival rate *ρ* combined with a smaller dilution frequency *λ* increases the chances of toxin-producers to win for all three policies. It is clear that the tactically-optimal (panel B) killers significantly outperform the constitutives (panel A). The differences in 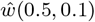 between *α*_∞_ and *α*_1_ are small, with the discrepancy increasing toward the upper right corner (panel C).

### Is the outcome affected by dilution strength and frequency?

The competition dynamics described so far appear to be robust, holding true for a variety of stochastic environments. In Fig. 8, we focus on a single initial condition (*f*_0_, *N*_0_) = (0.5, 0.1) and compare the performance of constitutive, tactically-optimal and strategically-optimal toxin-production policies for a range of (*ρ, λ*) values. Predictably, all three of them yield higher chances of winning against the toxin-sensitive strain when the dilutions are rare and weak (small *λ*, large *ρ*) – in these regimes, the population gets closer to the carrying capacity in between dilutions, the growth of both strains slows down, and the toxin’s effect becomes more noticeable. As expected, the constitutives are far less effective on most of this map. The biggest surprise is how well the tactically-optimal killers do – their chances of winning are in the worst case only 2.5% below those of the strategically-optimal killers. This is impressive since the tactically-optimal policy *α*_1_ is formulated without any reference to *ρ*, but works well across a fairly broad range of dilution strengths. But even against the strategically optimized *α*_∞_, the toxin-sensitives still have a chance of winning above 50% on at least half of this (*ρ, λ*) map.

In conclusion, whether dilutions are randomly timed or periodic and despite the benefits brought by toxin-production regulation to killers, the sensitive strain can prevail by taking advantage of disruptions caused by dilutions.

## Discussion

The impact of environmental variability on population size and diversity is well-established in the literature. For example, it is well known that *environmental switches* can make co-existence possible even if all the “un-switched” versions of the environment would result in a competitive exclusion with the same winner [69]. Similarly, the variability in the resulting population is influenced not only by the frequency but also by the timing (periodic vs random) of switching events [70]. While most studies employ environmental switches to model abrupt changes in population dynamics (e.g., instantaneous reductions in carrying capacity or resource processing efficiency), there is also a growing interest in understanding instantaneous exogenous events that impact all subpopulations similarly without affecting the dynamics in between. Dilutions provide a prime example of this, resulting in boom and bust cycles, where the bust phase is nearly instantaneous [71, 72]. Within this emerging focus, our paper is the first demonstration that for antagonistic interactions such population busts can fundamentally alter outcomes, determining which species achieves competitive exclusion.

We have shown both theoretically and experimentally that dilution events can benefit toxin-sensitive strains, when their rate of growth in the exponential phase is larger than that of toxin-producing ones. Using experiments with strains of *S. cerevisiae*, one engineered to constitutively produce the killer toxin K1, and the other sensitive to it, we found that the outcome and dynamics of competition between the two varied with the frequency of periodic dilution events. Because toxin production is often regulated in response to quorum sensing [54, 55], we used tools of optimal control theory to explore how different toxin-production policies can benefit the producers. Following this approach, we developed two efficient algorithms that (i) calculate the deterministic limit of the relative abundances of these strains as the number of periodic dilutions approaches infinity; and (ii) address the non-local Hamilton-Jacobi-type equation that includes the jumps introduced by dilution events in a stochastic environment. Our numerical experiments consistently support the conclusion that, regardless of toxin production regulation, dilution events often disrupt the dominance of the killer, thereby protecting the sensitive strain from extinction.

Rather than focusing on specific mechanistic models of toxin production regulated by quorum sensing, which would vary across species and would require a large number of parameters and modeling assumptions, we adopted a phenomenological approach to identify theoretical performance bounds for “omniscient killers” capable of measuring population density and fractions. The optimal policies derived here should thus be regarded as upper bounds on the ability of toxin-producing strains to out-compete sensitive ones in fluctuating environments. Future work will explore how more realistic, mechanistic models of toxin production regulated by quorum sensing compare to the optimal policies investigated here. For example, it will be useful to consider models in which the concentration of toxin and/or the concentration of quorum sensing molecules are additional state variables.

In a seminal paper on allelopathy in spatially distributed populations [25], Durrett and Levin showed that the competition of toxin-producing and toxin-sensitive strains displays bi-stability in well-mixed, undisturbed competitions, with either the killer or sensitive strain dominating in the long-term limit depending on the initial condition. In their model, such bi-stability depends critically on the magnitude of a (toxin unrelated) death rate term in their governing equations. In the SI Appendix §S2, we show that using typical laboratory-measured values for growth and death rates, the range of initial conditions for which the sensitive strain competitively excludes the killer is extremely small even under the Durrett-Levin model. This theoretical bi-stability alone is thus unlikely to explain why non-antagonistic strains are found in many microbial communities, and our analysis of disruption effects highlights another contributing factor, which is arguably at least as important. It is of interest to ask how the perturbations investigated here would affect the spatial competition between killer and sensitive cells explored theoretically in [25], where the two strain types were shown to coexist by forming dynamic, single-strain clusters. Similarly, it would be interesting to explore the impact of dilutions on the nucleation criteria that control the killers’ invasion success of a spatially-distributed, resident population [21].

As in any model, simplifying assumptions were made to focus on the main observed phenomena and to ensure the computational efficiency of our approach. But in the future, it might be desirable to relax some of these assumptions to reflect additional features present in realistic environments. One such extension would be to consider more general dilutions, for example, modeling survival rates *ρ* using a Binomial distribution, rather than assuming fixed relative abundances. Although Monte Carlo simulations (SI Appendix §S9) suggest that results remain qualitatively similar, random dilution outcomes can sometimes produce variability in which strain dominates, as seen in our experiments (Fig. 2C-D). An even more realistic model could also incorporate random variability in parameters such as growth rates (*r*_K_, *r*_S_), killing rates *γ*, and dilution factors *ρ* and *λ*, which are influenced by environmental and biological fluctuations [73, 74]. These fluctuations may better capture population dynamics but would increase computational complexity, requiring a hybrid model with discrete and continuous random perturbations based on jump-diffusion processes [75, 76].

Incorporating evolutionary adaptation, such as mutational dynamics, would also enhance our understanding of the competition between toxin-producing and toxin-sensitive strains in natural environments over longer timescales. Experiments indicate that killer strains can lose or alter their toxin-producing ability, and sensitive strains may develop resistance to the toxin [21, 23, 77]. The ultimate success of the killer strain in our model is negatively impacted by the costs associated with toxin production, suggesting that evolutionary adaptation may aim to minimize these costs [77]. Additionally, evolutionary adaptation may enable antagonistic strains to regulate toxin production in response to their environment, population abundance, or the presence of competitors [55], possibly approaching the performance of the optimal policies described here. Toxin resistance can arise through various mechanisms, such as alterations in toxin receptors or translocation pathways, which may have antagonistic pleiotropic effects where resistance incurs a cost in terms of growth rate [78]. This growth rate penalty will, in turn, influence the competitive dynamics with the killer strain. Finally, in environments experiencing disturbances, evolutionary adaptation may promote increased retention (*ρ*) [79] or even alter the rate of disturbances (*λ*). For example, production of surface-attachment molecules, pili or fimbriae by microbes such as *Pseudomonas aeruginosa, Vibrio cholerae, Clostridium difficile* and *Streptococcus salivarius* can help them adhere to surfaces in their environment and prevent them from being washed away in fluid environments such as the gastrointestinal tract, the oral cavity, or natural water bodies [11, 80, 81]. *C. difficile* and *V. cholerae*, in addition to adhering to surfaces such as the intestinal mucosa, can also cause diarrhea [82] and thus potentially control the environment dilution rate and intensity, at least transiently.

In conclusion, we posit that the fitness cost incurred by toxin-producing strains may be particularly detrimental in boom-and-bust environments in which populations undergo regular or stochastic dilutions. We propose this as a possible explanation for why both antagonistic and non-antagonistic microbes are found in nature, and why environments with higher turnover rates may favor the latter [51, 52].

## Materials and Methods

### Competition experiments

The killer strain yAG171b expressed the toxin gene K1 from the chromosome from the galactose-inducible *P*_*GAL*1_ promoter. To enable strain differentiation at the flow cytometer, strain yAG171b expressed the fluorescent protein CyOFP1opt [83], whereas strain yAG177 expressed the fluorescent protein ymCitrine (Table 3). Cells of the two strains could be clearly identified via flow cytometry and separated in the FL-1A (533 ± 30 nm) vs FL-2A (585 ± 40 nm) emission channels with excitation at 488 nm. Dead yAG177 cells (confirmed via Propidium-Iodide staining) formed a separate cluster in the FL-1A vs FL-2A scatter plot, and were excluded when calculating killer fractions. No dead cells were observed when growing killer cells in isolation. To prevent metabolization of galactose and to make the expression of K1 titratable, both *GAL1* and *GAL10* were deleted, and *GAL*3 was placed under the constitutive promoter *P*_*ACT*1_ [84], in both strains. In addition, to prevent catabolite repression that would have prevented expression of the K1 toxin from *P*_*GAL*1_ in the presence of glucose, we deleted *HXK2* in yAG171b. This deletion confers a growth-rate deficit to yAG171b, compared to yAG177, in addition to the cost of toxin production. Strain construction is described in the SI Appendix §S1.2. Experiments were performed with strains yAG171b (killer) and yAG177 (sensitive) in filter-sterilized Complete Synthetic Medium (CSM) buffered at pH 4.5 with succinic acid. The medium composition is described in the SI Appendix §S1.2. The medium was supplemented with 500 uM galactose for induction of the killer toxin gene K1. Experiments were performed in 96 well plates with flat bottom and a transparent lid, incubated in a plate reader at 23°C with orbital shaking at 425 rpm with an amplitude of 3 mm. For the experiment of Fig. 1, we initialized six technical replicates for each initial fraction and initial population size. Initial fractions of killers were measured via flow cytometry. The experiment was initialized with overnights grown in CSM at 30°C, which were diluted in CSM with 500 *µ*M galactose by a factor 3125 (*N*_0_ treatment), 312.5 (10 ·*N*_0_ treatment), 156.25 (20· *N*_0_ treatment) and 62.5 (50 · *N*_0_ treatment). For the experiments of Fig. 2, we initialized four technical replicates for six different 24 h and 48 h dilution cycle treatments, targeting initial killer fractions of 32%, 36%, 40%, 44%, 48% and 52%. Actual initial fractions were measured via flow cytometry and were within ±1% of the targeted ones; see Fig. 2. The experiment was initialized with overnights grown in CSM at 30°C, which were diluted by a factor 3125 in CSM with 500 *µ*M galactose. Successive daily dilutions were performed with a dilution factor of 225. In both experiments, the culture volume was 150 *µ*L per replicate.

## Acknowledgments

We thank Andrew W. Murray, Stephen P. Ellner, and Tobias Dörr for insightful comments on the manuscript. AG thanks Marie F. Gorwa-Grauslund for gifting a plasmid with *CyOFP1opt*. AG acknowledges support by the National Institute of General Medical Sciences of the National Institutes of Health, United States of America under award number 1R35GM147493. MW and AV acknowledge support by the National Science Foundation (awards DMS-1645643 and DMS-2111522) as well as the Air Force Office of Scientific Research (award FA9550-22-1-0528).

## Author Declaration

The authors declare no conflict of interest.

## Supporting Information (SI) Appendix

SI Appendix provides additional details for the main text. We continue to use reference numbers of equations and figures previously introduced in the main text. The numbering of figures in SI text is prefixed with an “S”, and bibliographic references used here are listed and numbered separately.

## S1 Experiments

### S1.1 Additional experimental results

Fig. S1 shows cell density estimates obtained from Optical Density (OD600) measurements taken throughout the experiment of Fig. 2, showing that populations diluted every 24 h (teal curves) did not reach carrying capacity, whereas those diluted every 48 h (pink curves) did.

**Figure S1.**
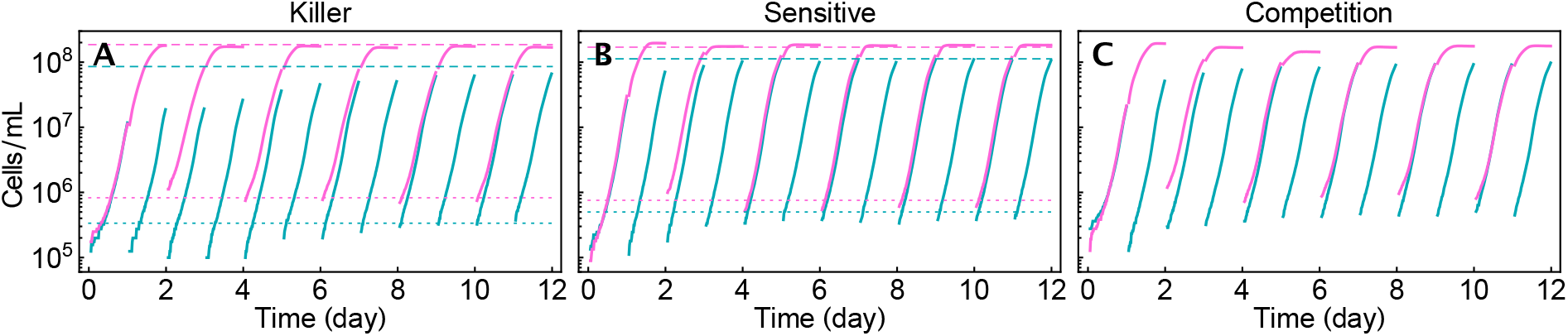
Mean cell density values recorded throughout the experiment of Fig. 2. The mean cell densities in the 24 h cycle replicates are shown in teal, and the 48 h cycle ones are in pink. (A) Mean cell density across three replicates initialized with 100% killer. (B) Mean cell density across three replicates initialized with 0% killer. (C) Mean cell density across four replicates initialized with 44% killer. Cell densities were estimated from Optical Density (OD600) data via a cubic calibration curve. Dashed and dotted lines are predictions of asymptotic pre- and post-dilution cell densities predictions based on Eq. (S3.1.3) with non-normalized population sizes.

### S1.2 Strains, oligos, plasmids and additional methods

The Complete Synthetic Medium (CSM) composition was, per liter: 0.59 g of amino acid mix, 1.706 g of Yeast Nitrogen Base without amino acids, carbohydrates, ammonium sulfate, ferric chloride and copper sulfate (USBiological Y2030-03), 20 g of dextrose, 4 mL of a 5 g/L L-histidine hydrochloride monohydrate stock solution, 20 mL of a 5 g/L L-leucine stock solution, 8 mL of a 2.5 g/L uracil stock solution, 20 mL of a stock solution containing 5 g/L adenine hydrochloride and 5 g/L L-tryptophan, 11.2 g of succinic acid, 5 g of ammonium sulfate and NaOH to reach pH 4.5. The amino acid mix was made by mixing 10 g L-arginine hydrochloride, 16 g L-aspartic acid, 10 g L-isoleucine, 10 g L-lysine hydrochloride, 4 g L-methionine, 10 g L-phenylalanine, 20 g L-threonine, 10 g L-tyrosine and 28 g L-valine.

Tables 1, 2, and 3 report oligos, plasmids and strains used for the experiments. We substituted ymCherry in pAG11 with CyOFP1opt from pRP008 using Gibson assembly, producing pAG134, whose sequence was verified via long-read sequencing. Strain yAG171 was obtained by digesting pAG134 with PpuMI and transforming it into yAG75 [1] for integration of pAG134 in T_CYC1_. Strain yAG177 was obtained by digesting pAG5 [1] with PpuMI and transforming it into yAG74 [1] for integration of pAG5 in P_ACT1_.

**Table 1:**
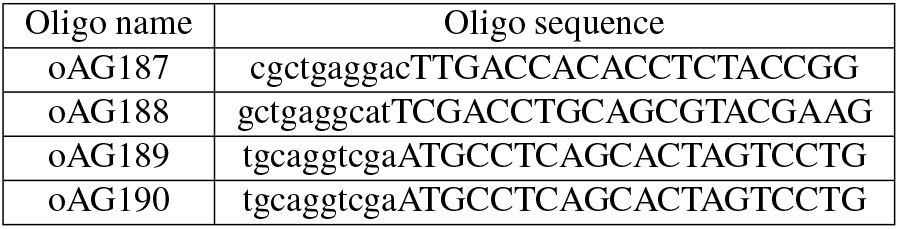
DNA oligos used to assemble pAG134 via Gibson assembly. Bases in capital letters represent homology to the PCR template (pAG11 for yAG187/188 and pRP008 for yAG189/190) Lowercase bases represent homology to the backbone or fragment for Gibson assembly.

**Table 2:**
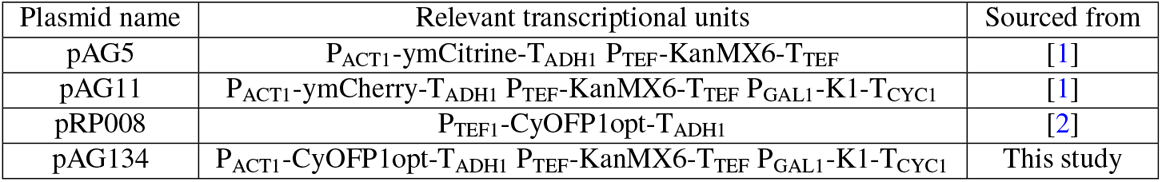
Plasmids used for strain construction.

**Table 3:**
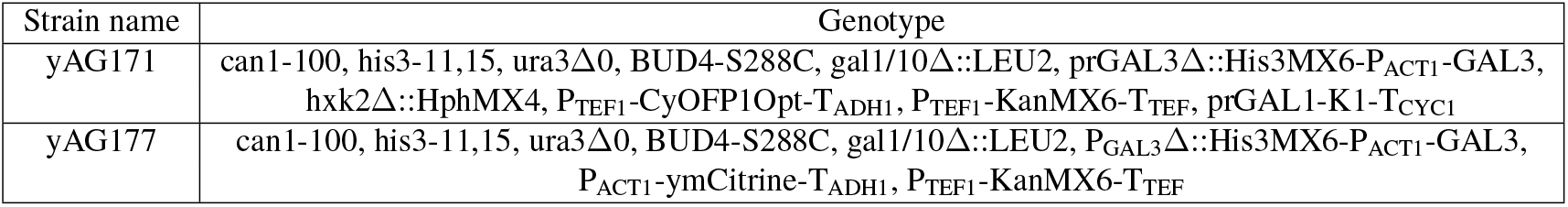
Strains used for the experiments.

## S2 Comparison with the Durrett-Levin model

In the main text, all of our results are based on a basic competition model between the toxin-producing “killer” strain (K) and the toxin-sensitive strain (S) (Eq. (1) of the main text). This model assumes the growth of both strains is logistic with respective intrinsic growth rates (*r*_K_, *r*_S_) and a shared carrying capacity *C*. The toxin-induced death rate of the sensitive cells is proportional to the product of the strain densities, *n*_K_*n*_S_. Focusing on *constitutive killers* who produce the toxin at the maximal rate *a* = 1, the original Eq. (1) reads

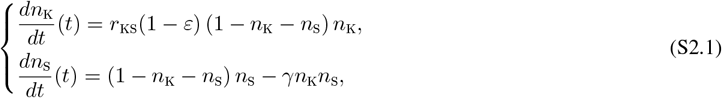

where *r*_KS_ = *r*_K_/*r*_S_< 1 is the intrinsic growth rates ratio, *γ* is the killing rate rescaled by *r*_S_, and *ε* is the cost associated with producing toxin at the maximal rate (*a* = 1). We note three equilibria (fixed points) of Eq. (S2.1):

1. (*n*_K_, *n*_S_) = (0, 0) – a nodal source
2. (*n*_K_, *n*_S_) = (1, 0) – a nodal sink
3. (*n*_K_, *n*_S_) = (0, 1) – a degenerate node

It follows that a competitive exclusion of the sensitive by the killer will be observed starting from any non-equilibrium initial condition. Despite this eventual dominance, the sensitive strain may transiently increase its relative abundance depending on the initial configuration; see Fig. S2. This occurs because the growth rate benefit of not producing the toxin initially outweighs the deficit caused by toxin-induced death. Fig. S3D further illustrates this phenomenon in a phase plane, showing that the sensitive population begins to decline once the total population reaches a sufficiently large size.

**Figure S2.**
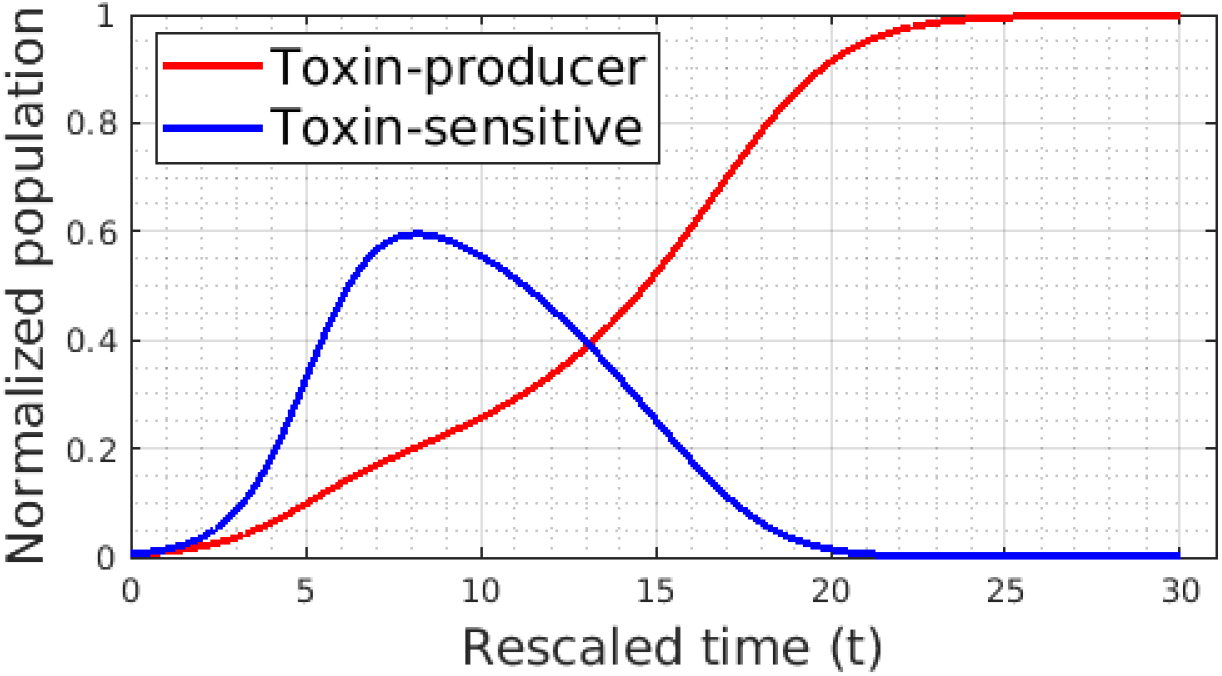
Population trajectories for a constitutive toxin-producing strain (based on Eq. (S2.1)) competing against a toxin-sensitive one. Starting with 50% of each strain and a total population equal to 1% of the carrying capacity, sensitives initially grow faster due to their higher intrinsic growth rate. However, as the population size approaches the carrying capacity, the intrinsic growth advantage of the sensitive strain diminishes, leading to eventual domination by the toxin producer due to the toxin’s action. Parameter values: *r*_KS_ = 0.85, *ε* = 0.2, and *γ* = 1.

One natural extension of model Eq. (S2.1) is to incorporate natural (toxin-unrelated) death rates (*δ*_K_, *δ*_S_) for both strains. With both of these rates rescaled by 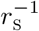, this yields a system of ODEs equivalent to a competition model previously studied by Durrett and Levin [3]:

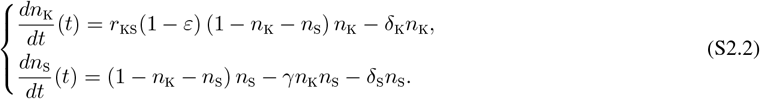

By setting both equations to zero, we find four fixed points of this system:

1. (*n*_K_, *n*_S_) = (0, 0)
2. 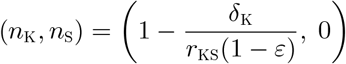
3. (*n*_K_, *n*_S_) = (0, 1 − *δ*_S_)
4. 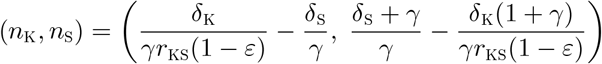

Stability analysis shows that the origin is a *nodal source*, the two boundary equilibria are both *nodal sinks*, and the interior fixed point is a *hyperbolic saddle* if

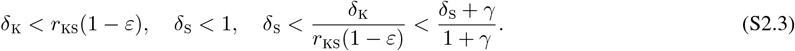

For this range of parameters, the system exhibits “bi-stability”: starting above the saddle’s stable manifold (the magenta dash-dotted line in Fig. S3A) leads to competitive exclusion of the killer by the sensitive, while starting below it results in competitive exclusion of the sensitive by the killer. See Fig. S3A for a detailed phase portrait. It is worth noting that, when the natural death rates approach zero (i.e., as *δ*_S_, *δ*_K_ → 0), Eq. (S2.2) reduces to Eq. (S2.1). In the meantime,

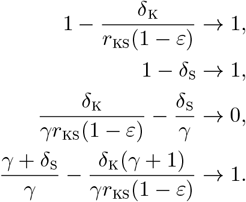

Consequently, the last two equilibria collapse to (*n*_K_, *n*_S_) = (0, 1) while the second fixed point moves to (*n*_K_, *n*_S_) = (1, 0). This means the system loses its “bi-stability” structure as the “sensitive-winning” region shrinks when the natural death rates approach zero (see the transitions in Fig. S3). Schink et al. [4] estimated the death rate of an *E. coli* strain K-12 at 0.018 h^−1^ in laboratory experiments, whereas the max growth rate was 0.7 h^−1^. In our time scale, this corresponds to *δ*_K_ = *δ*_S_ ≈ 0.0257. We see from Fig. S3C that in this scenario, the hyperbolic saddle is extremely close to the “all sensitives” equilibrium and both of them are close to (*n*_K_, *n*_S_) = (0, 1). This extremely small basin of attraction for the “all sensitives” equilibrium makes it harder to explain why sensitive strains are often found in the natural environment. This issue is one of the motivations for the current paper, and for our conjecture that, with dilutions, the sensitives can win even in the *δ*_K_ = *δ*_S_ = 0 limit, where the bi-stability disappears; Fig. S3D.

**Figure S3.**
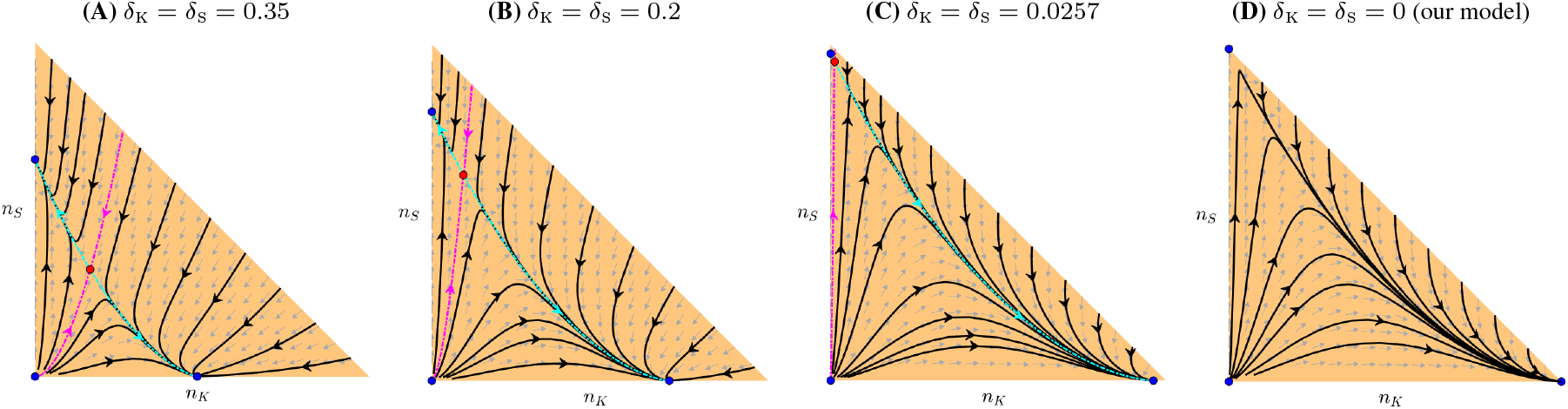
Phase portraits of the Durrett-Levin model (S2.2) with decreasing death rates. The “bi-stability” is noticeable when the death rates (*δ*_K_, *δ*_S_) are comparable in magnitude to the growth rates (panels (A,B)). Using laboratory-estimated death rates values [4], the hyperbolic saddle moves toward the “all sensitives” stable node, with both converging toward (*n*_K_, *n*_S_) = (1, 0) (panel C). When (*δ*_K_, *δ*_S_) = (0, 0), the Durrett-Levin model reduces to our model (S2.1), and “bi-stability” disappears (panel D). In (A-C), the hyperbolic saddle is plotted with a red dot while other equilibria are plotted with a blue dot. The stable manifold is plotted with a magenta dotted-dashed line while the unstable manifold is plotted with a cyan dotted-dashed line. In (D), all equilibria are plotted with a blue dot. In all of them, gray arrows denote the vector field directions corresponding to Eq. (S2.2). Parameter values: *r*_KS_ = 0.85, *ε* = 0.2, and *γ* = 1.

## S3 Effect of dilutions on a single-strain logistic growth model

This section presents theoretical and numerical results relevant for a *single* strain subjected to either regular or randomly timed dilutions. These findings provide background information for the main text and are relevant after one of the strains becomes strongly dominant.

### S3.1 Effect of regular dilutions

In this subsection, we prove the theoretical pre-dilution population limit for a population growing according to the logistic model and undergoing regular dilutions.

#### Theorem S3.1.

*Consider the following rescaled logistic growth model*

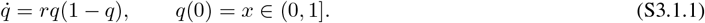

*If the system undergoes regular dilutions with a fundamental period of T, and after each dilution, a deterministic fraction ρ of the population survives, i.e*.,

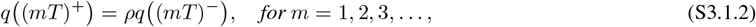

*then*

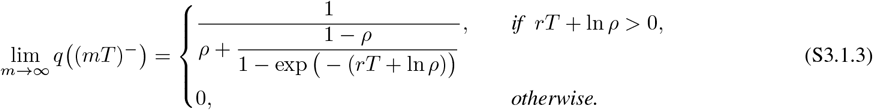

*Proof*. We begin the proof of Theorem S3.1 by proving the following lemma.

#### Lemma S3.2.

*Given the rescaled logistic model Eq. (S3.1.1), the pre-dilution population size up to the m-th cycle is*

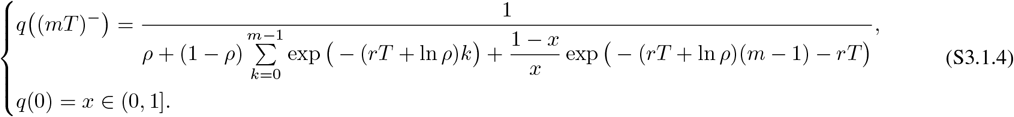

*Proof*. We prove the above lemma by mathematical induction. Starting with the first dilution (*m* = 1), from Eq. (S3.1.4) we have

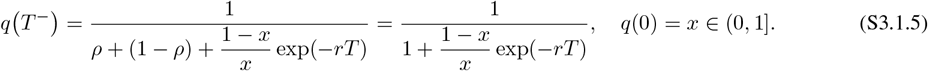

We show it is true by solving Eq. (S3.1.1) analytically. By separation of variables, one can show that the general solution to Eq. (S3.1.1) is

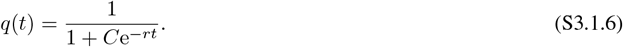

With *q*(0) = *x*, we have 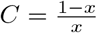. Consequently,

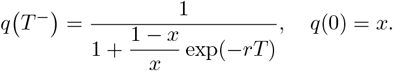

Now assume Eq. (S3.1.4) holds true for some integer *j* > 1. I.e.,

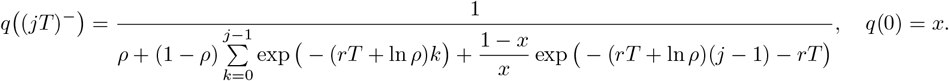

We now show it holds for *j* + 1. Notice that *q*([(*j* + 1)*T* ]^−^) with *q*(0) = *x* is equivalent to *q*(*T*^−^) with *q*(0) = *q*((*jT*)^+^) =*ρq*((*jT*)^−^) under the assumption of proportional dilutions. Given the general solution to Eq. (S3.1.1) in Eq. (S3.1.6), we compute the arbitrary constant *C* by imposing the new initial condition *q*(0) = *ρq*((*jT*)^−^). It follows that

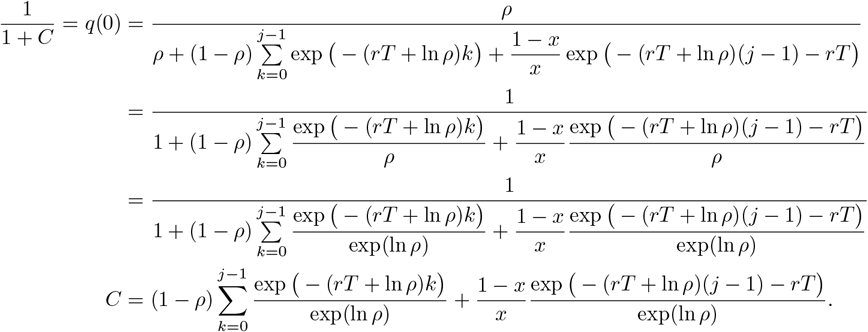

Substituting it back into Eq. (S3.1.6), we have

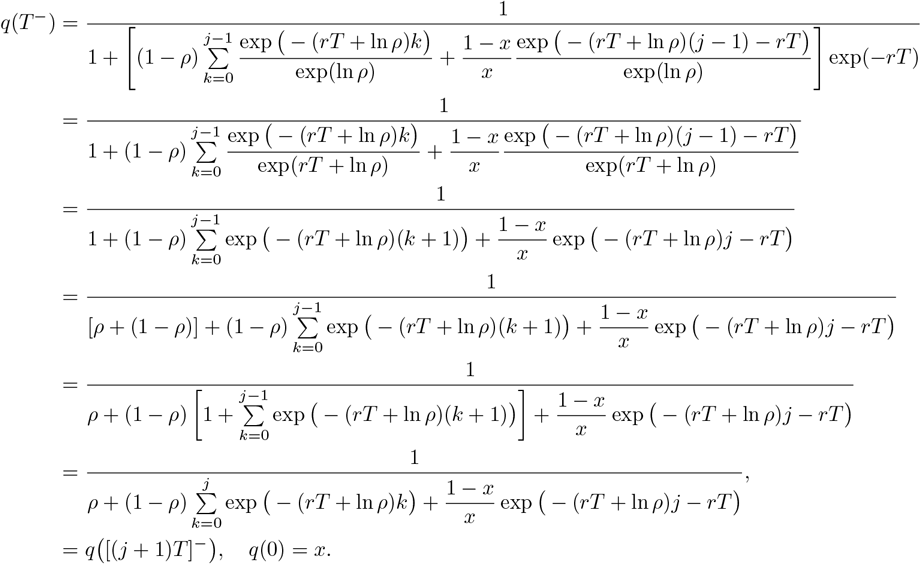

Consequently, Eq. (S3.2) holds for all *j* ∈ *ℕ*.

Now, we take the limit of Eq. (S3.2) as *m* approaches infinity. Notice that the second term in the denominator is a geometric series. It follows that

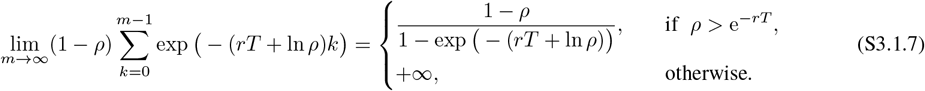

With the third term in the denominator of Eq. (S3.1.4) approaching 0 as *m* → ∞, we thus conclude that

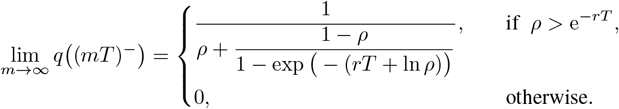

Therefore, for a population consisting of sensitives only, i.e.,

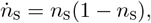

the limiting pre-dilution population size is

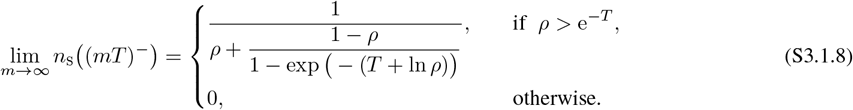

For a population consisting of toxin-producing killers only, i.e.,

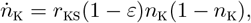

the limiting pre-dilution population size is

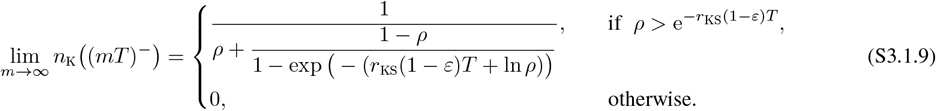

Fig. S4 shows the numerical values derived in Eq. (S3.1.8) and Eq. (S3.1.9) in the (*ρ, T*) phase plane. It is evident that the larger the proportion of the population surviving after dilution (higher *ρ*) and the longer the period (higher *T*), the larger the pre-dilution population size as the number of dilutions approaches infinity. For non-zero limits, sensitives can maintain a higher limiting population than any killers due to their faster reproduction rate.

**Figure S4.**
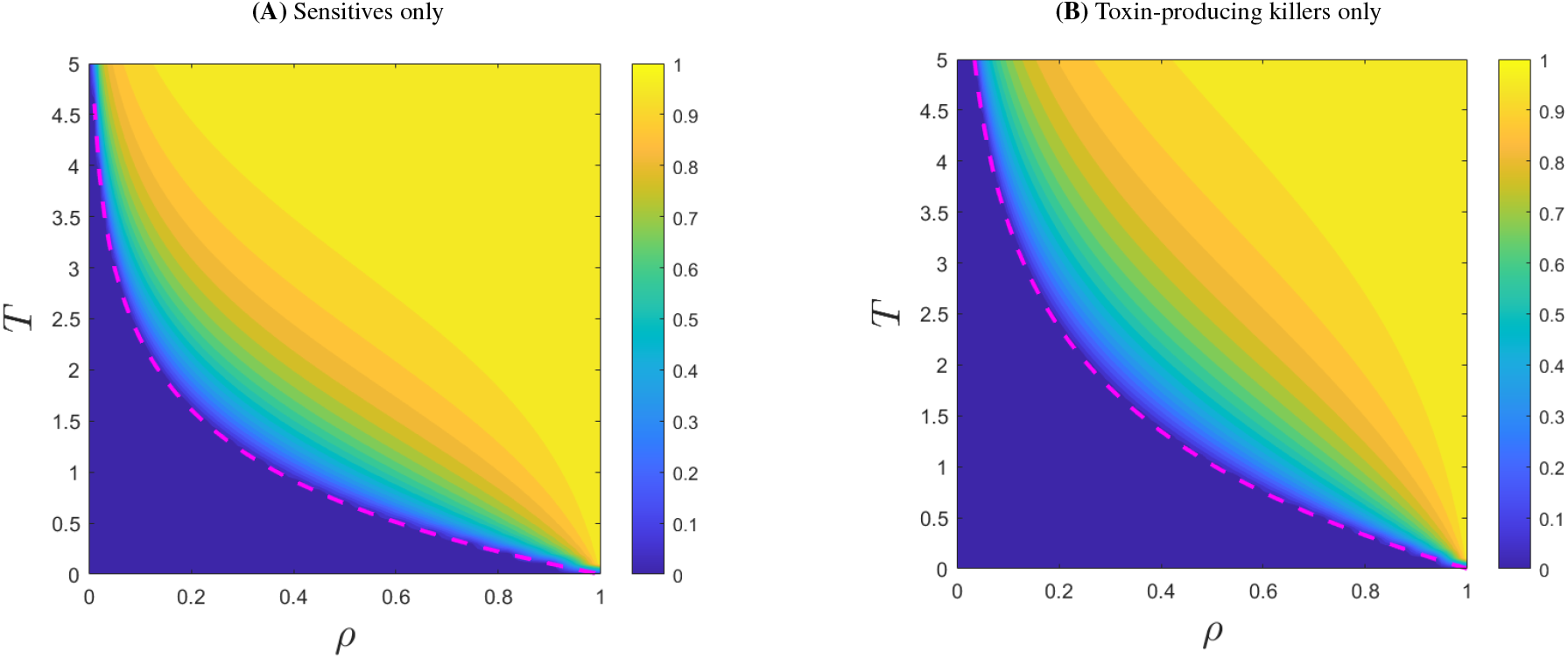
Pre-dilution single population limits under regular dilutions in (*ρ, T*) phase plane. (A) sensitives only (*r* = 1); (B) Toxin-producing killers only (*r* = *r*_KS_(1 − *ε*) = 0.68). In both of them, the magenta-dashed line corresponds to *ρ* = exp(−*rT*) with their respective intrinsic growth rate *r*. The limiting pre-dilution population is zero below this line.

### S3.2 Effect of randomly-timed dilutions

Next, we consider randomly timed dilutions, following a Poisson process with rate *λ*. In this case, any inter-dilution time T is exponentially distributed with the expected value E[T] = 1/*λ*. To compare with the previous results, we focus on the *expected pre-dilution population size* as the number of dilutions approaches infinity, with the population growing according to Eq. (S3.1.1). We still assume that a fixed fraction *ρ* of the population survives after each dilution. As an analytical limit is unlikely to be obtained, we conducted Monte Carlo simulations with 200 dilutions starting from the initial population *q*(0) = 0.5. Fig. S5 shows that the general trend remains unchanged: weaker dilution strength (higher *ρ*) and slower arrival rate (lower *λ*) result in higher expected pre-dilution population size in the limit. However, with randomly timed dilutions, the region where the averaged population is nearly 1 in the limit is significantly reduced in both cases. Surprisingly, the previous “deterministic boundary” *ρ* = exp(−*r/λ*) (the magenta dashed line in Fig. S5) still appears to accurately predict where the population goes extinct in the limit in the (*ρ*, 1/*λ*) phase plane.

**Figure S5.**
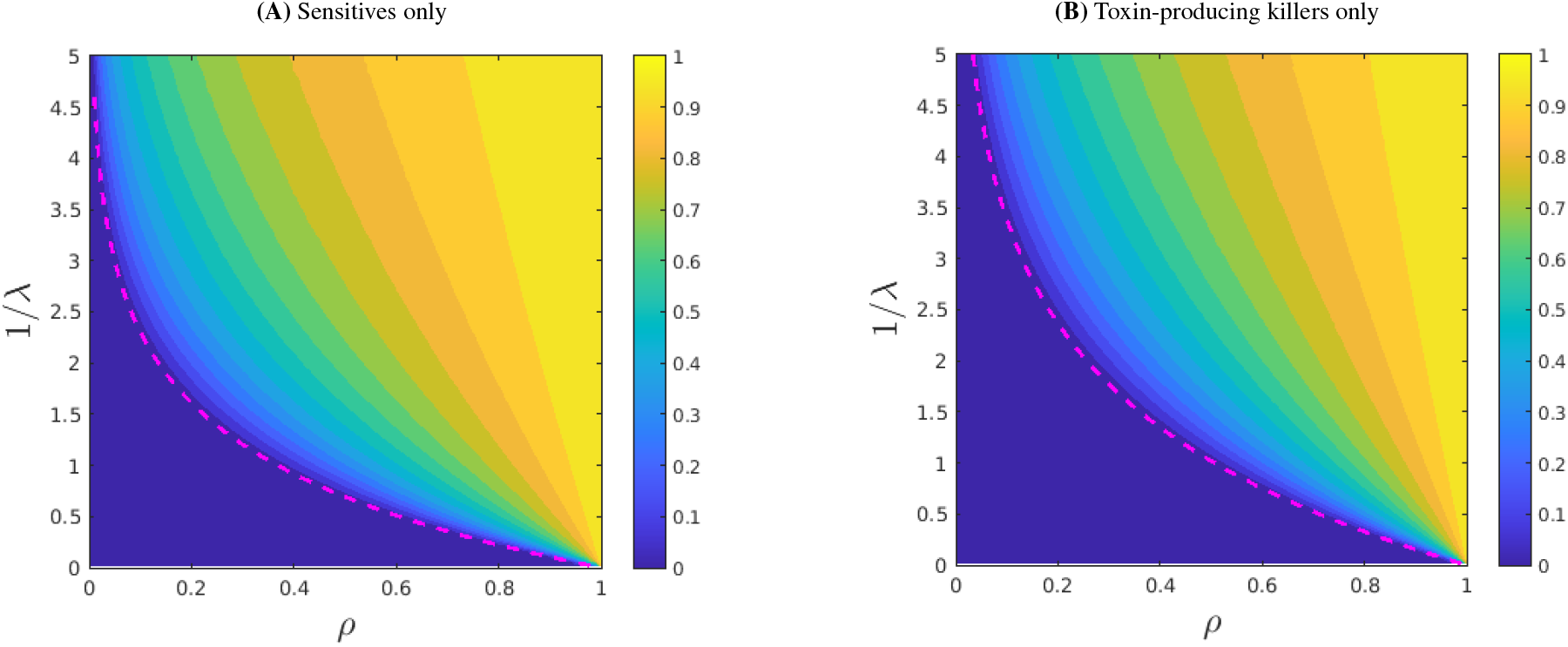
Randomly-timed dilutions: mean empirical population just before the 201st dilution shown in the (*ρ*, 1/*λ*) phase plane. (A) sensitives only (*r* = 1); (B) Toxin-producing killers only (*r* = *r*_KS_(1 − *ε*) = 0.68). In both panels, the magenta-dashed line corresponds to *ρ* = exp(−*r/λ*) with their respective intrinsic growth rate *r*. The population below this line will most likely go extinct. Both panels are produced by Monte Carlo simulations on a uniform grid with gridpoints (*ρ*_*i*_, (1/*λ*)_*j*_) = (*i*/100, 0.01 + *j*/20) where *i, j* = 0,…, 100. At each gridpoint (representing a particular environment), we used 10^5^ random population trajectories starting from the same initial population size of 0.5.

## S4 Who benefits from randomness in dilution times?

To check whether the randomness in dilution times and the values of the surviving fraction *ρ* and dilution frequency *λ* affect the performance of constitutive and myopic/tactically-optimal toxin production policies, we introduce a new metric of competitive advantage and use it across a range of (*ρ, λ*) values. Assuming that the initial population size *N*_0_ is fixed while *f*_0_ ∈ [0, 1] is selected uniformly at random, we examine the probability of killers’ winning. For regular/deterministically-timed dilutions with period *T* = 1/*λ*, this probability (denoted by *L*(*N*_0_)) is simply the width of the killer’s “deterministically-winning” (red) region on the horizontal line *N* = *N*_0_, e.g., in Fig. 4C-D. With random dilution times, this probability is 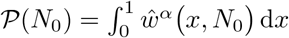. To quantify the impact of randomness, we define 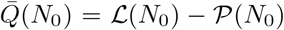. Focusing on *N*_0_ = 0.5, Fig. S6 presents a heat map of 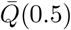 for various (*ρ, λ*) values. Panel A shows that, for constitutive killers, the randomness is beneficial in the upper left half of the parameter space (where dilutions are more severe and frequent) but actually slightly detrimental in the bottom right half (where dilutions are more moderate and rare). In contrast, for the population-sensing (myopic, optimized for a specific *T* or *λ*) killers, the randomness in dilution times appears to be beneficial across all (*ρ, λ*) values.

**Figure S6.**
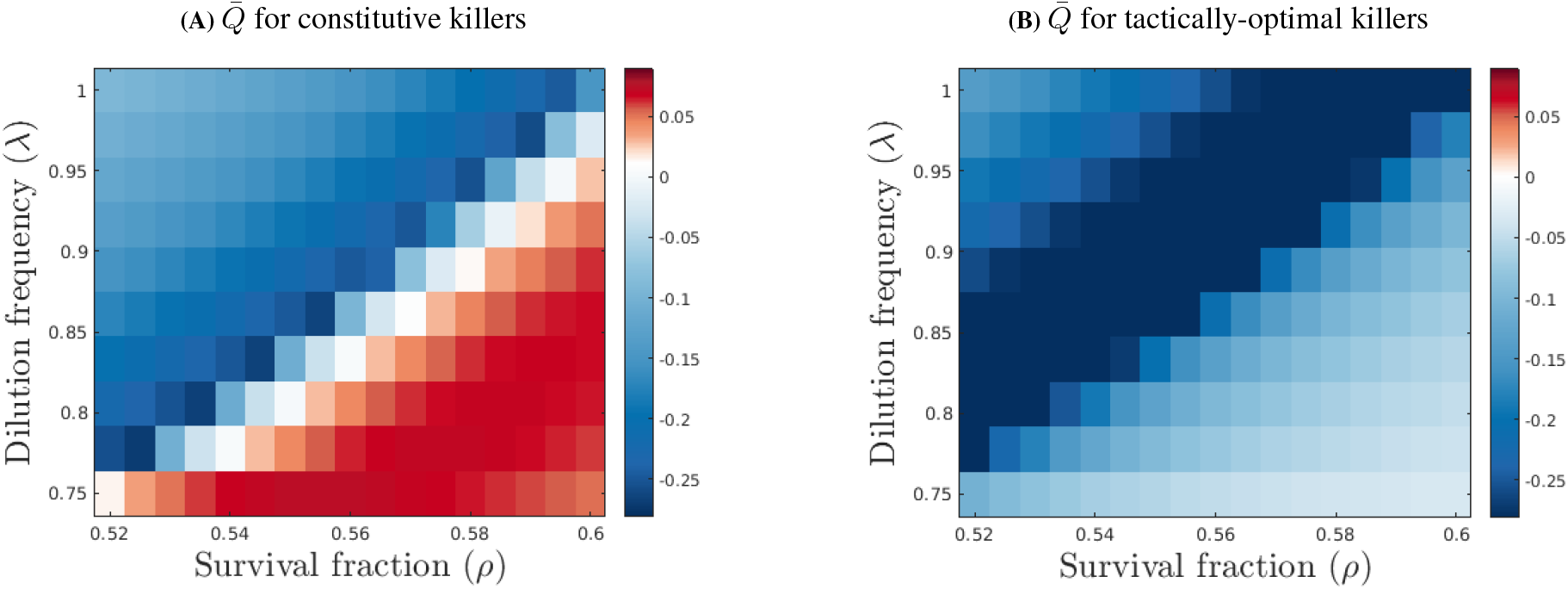
The impact of dilution-time randomness and intensity on the performance of constitutive and tactically-optimal killers across a range of (*ρ, λ*) values. The performance metric 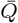 (defined in the text) is shown in red wherever the toxin-producers have better chances of winning with regular/periodic dilutions and in blue wherever their chances are better with randomly-timed dilutions (assuming the same average frequency: *λ* = 1/*T*). (A) For constitutive killers, the randomness is beneficial when dilutions are more severe and frequent, but it is sightly detrimental when dilutions happen more rarely and are less drastic. (B) For tactically-optimal killers (with policies optimized for each *λ* and *T* = 1/*λ*), the randomness is beneficial across all tested parameters.

## S5 Comparison of strategically and tactically-optimal killers and constitutive ones

Fig. S7B shows that the strategically-optimal policy *α*_∞_ prescribes producing toxin slightly more conservatively than the tactically-optimal *α*_1_, but in the end its performance is only marginally better for our chosen parameter values (Fig. S7D). However, both of these population-sensing-enabled policies have a very significant advantage over the constitutive *α*_0_ = 1; see Fig. S7C and Fig. 8 in the main text.

**Figure S7.**
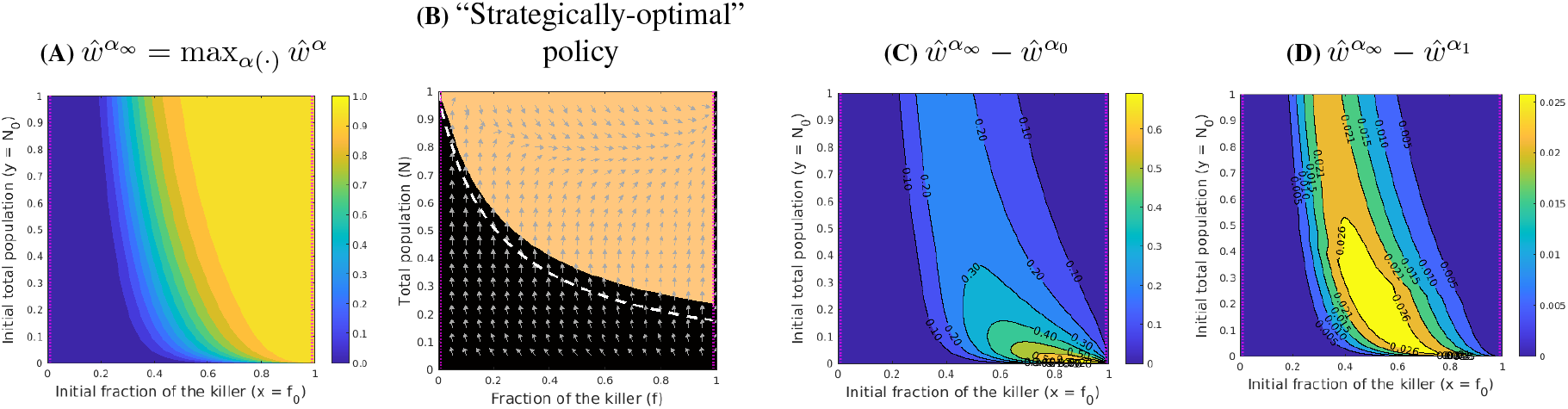
“Strategically-optimal” killers: performance (A), policy (B), and comparison with constitutive (C) and tactically-optimal (D) killers. The optimal toxin-on region for the “strategically-optimal” killers (orange in B) slightly shrinks compared to that of the “tactically-optimal” killers (the boundary of which is shown by a white-dashed line). As a result, the maximized probability of winning (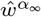in panel A) is only marginally better than 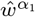, with a maximum difference of just 0.025 (see the absolute difference map in D). However, compared to constitutive killers, the advantage is significant: on a large part of the domain, the improvement in the chances of winning is above 20% (panel C). For really small *N* and relatively large *f*, this advantage is even above 60% – this is the set of initial conditions where constitutive killers grow much slower and are thus more affected by occasional short inter-dilutions intervals. In all panels, the victory and defeat barriers (*γ*_v_ and *γ*_d_, respectively) are plotted with a magenta dotted line. In (B), gray arrows denote the vector field directions corresponding to Eq. (2) in the main text with *a* = *α*_∞_ (*f, N*). In both (C) & (D), the contour lines are labeled with their respective probability values.

## S6 Derivation of Hamilton-Jacobi-Bellman equations

In this section, we derive the HJB equations for “tactically-optimal” killers (Eq. (11) in Box 4) and the “strategically-optimal” killers (Eq. (14) in Box 4) via tools of dynamic programming. For the former, see also [5, 6] for more details. The derivation of the time-dependent HJB PDE for the finite-horizon problem (Eq. (3) in the main text) is omitted here, as it can be easily found in classical literature, such as [7].

Recall that we model the random dilution events as a Poisson process with a fixed rate *λ* > 0. Thus, any inter-arrival time 𝒯 is exponentially distributed with rate *λ*. We assume that after each dilution, the fractions (relative abundances) are preserved while only a fraction *ρ* of the total population survives.

### S6.1 “Tactically-optimal” killers

Recall the value function for the “tactically-optimal” killer in Box 4 in the main text:

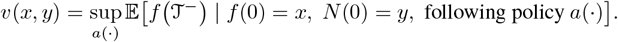

Since 𝒯 ∼ Exp(*λ*), the expectation is defined as

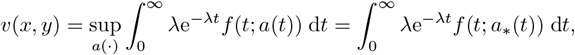

with an assumption that a maximizing/optimal policy *a*_*_(·) exists. Rewriting this formula as

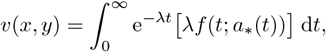

we can now re-interpret it as an *infinite horizon* problem with running cost *λf* (*t*; *a*_*_(*t*)) and a discounting factor *λ*.

For a sufficiently small *h>* 0, by Bellman’s Optimality Principle, we have

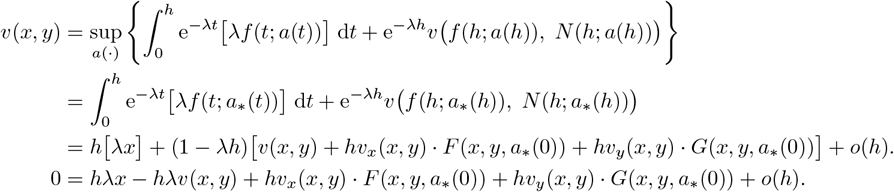

Now dividing both sides by *h* and sending *h* to 0, we obtain

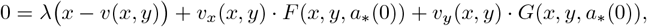

or, more generally,

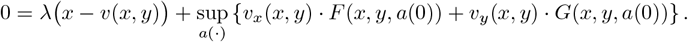

Notice that the above equation involves *a*_*_(0) only. It is then natural to switch to a state-dependent optimal control in feedback form. The HJB equation that *v* satisfies is then obtained by maximizing over *a* = *a*(0) ∈ [0, 1]. By demanding the above equation holds for all (*x, y*) ∈ [0, 1]^2^, the PDE can be written as:

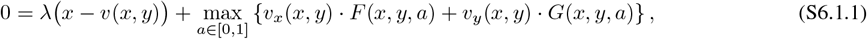

with the boundary condition

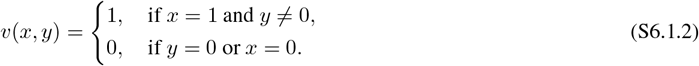

Substituting the actual definitions of *F* and *G*, we obtain Eq. (S6.1.1) in the specific form:

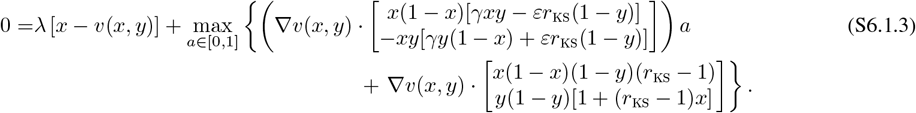

The linear dependence on *a* yields the *bang-bang* property:

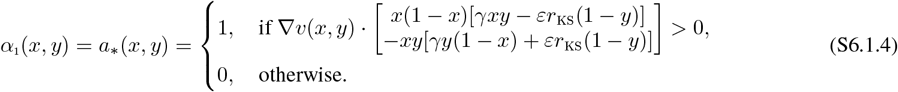

### S6.2 “Strategically-optimal” killers

Recall the corresponding value function Eq. (13) from the main text:

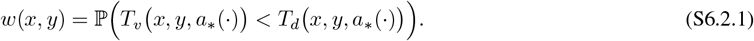

Let 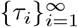 be an infinite sequence of random dilution times and *t*_*m*_ = *m*Δ*t, m* = 0, 1, 2, 3,… be a uniform time discretization. We again assume the optimal policy *a*_*_ exists, and let

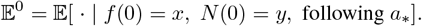

Thus, by Bellman’s Optimality Principle and the *law of total expectation*, we have

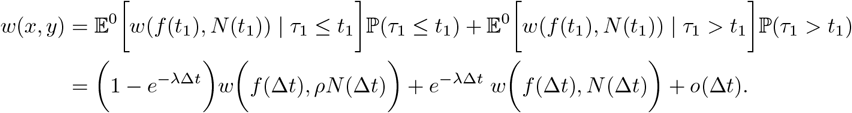

A first-order approximation around *t*_0_ = 0 gives

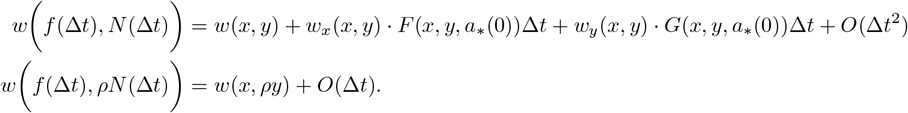

And hence

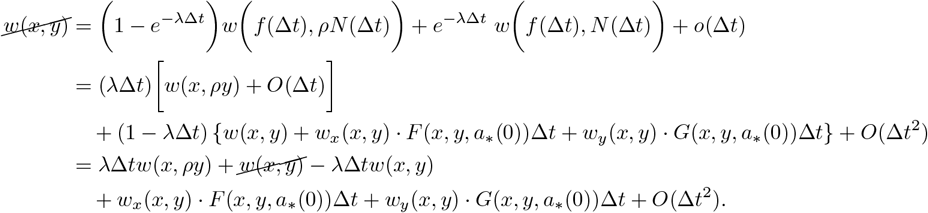

Dividing it by Δ*t* and taking Δ*t* ↓ 0, we have

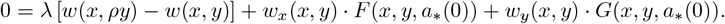

Notice that the above equation involves *a*_*_(0) only. It is then natural to switch to a state-dependent optimal control in feedback form. The HJB equation for Eq. (13) is then obtained by maximizing over *a* = *a*(0) ∈ [0, 1]. By demanding the above equation holds for all (*x, y*) ∈ [0, 1]^2^ \ Δ (where Δ is the terminal set, see Box 3), the PDE can be written as:

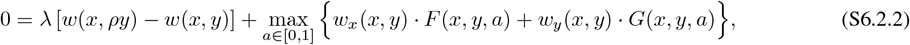

with the boundary condition

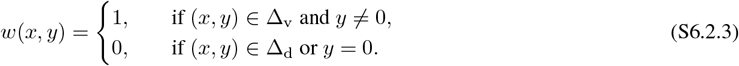

Substituting the actual definitions of *F* and *G*, we obtain Eq. (S6.2.2) in the specific form:

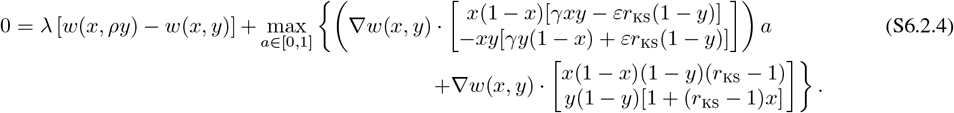

The linear dependence on *a* yields the *bang-bang* property:

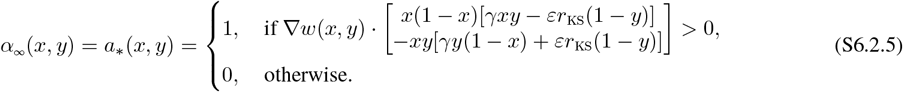

As mentioned in the main text, our system dynamics with randomly timed dilutions can be interpreted as a Piecewise-Deterministic Markov Process (PDMP). In general, the value function associated with a PDMP might not be smooth or even continuous. However, it can still often be interpreted as a unique (discontinuous) *viscosity solution* of the HJB equation [8].

#### Remark I

The linear non-local Eq. (9) in Box 3 can be derived in the same way but using a fixed policy 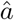 instead of *a*_*_.

#### Remark II

Let

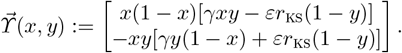

For both the “tactically-optimal” killer and the “strategically-optimal” killer, the respective HJB equation is linear in *a*, which yields a *generally* bang-bang optimal policy. However, we note that singular controls may arise when either 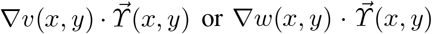 is equal to 0. But by computing the vector field associated with both *α*_1_ and *α*_∞_, we find no vector tangential to the boundary of *a*_*_(*x, y*) = 1, thereby excluding the possibility of singular arcs in our models. The same situation applies to the optimal policy *a*_*_(*x, y, t*) for the regular/period dilutions as well.

## S7 Numerical methods and implementation details

In this section, we provide the numerical schemes and implementation details of solving: (i) the finite horizon HJB PDE for *u*(*x, y, t*) with regular/periodic dilutions in §S7.1; (ii) the limiting fractions 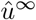 and total population 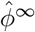 under regular dilutions in §S7.2; and (iii) the non-local HJB equation for *w*(*x, y*) for the “strategically-optimal” killers in §S7.3. For (i) and (iii), the optimal feedback policy is found by numerically solving the corresponding HJB equation.

### S7.1 For the finite-horizon HJB

Recall from the main text that we define the fraction-maximizing *value function* as

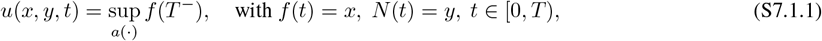

where *u* satisfies a time-dependent HJB PDE

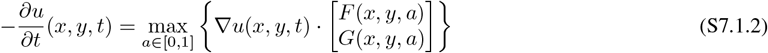

with the terminal condition

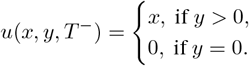

Substituting the actual definitions of *F* and *G*, we obtain Eq. (S7.1.2) in the specific form:

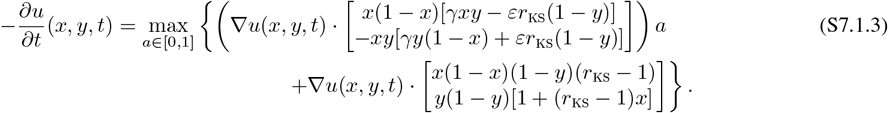

The linear dependence on *a* yields the *bang-bang* property:

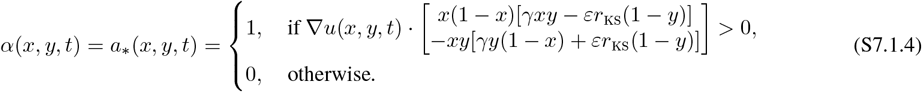

The time-dependent total population, denoted by *ϕ*(*x, y, t*), satisfies a linear PDE

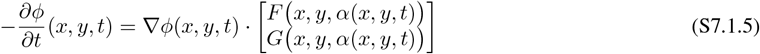

with the terminal condition *ϕ*(*x, y, T* ^−^) = *y*.

We approximate the solution to Eq. (S7.1.2) by a first-order semi-Lagrangian discretization [9] on a uniform rectangular grid over the (*x, y, t*) space. I.e., (*x*_*i*_, *y*_*j*_, *t*_*k*_) = (*i*Δ*x, j*Δ*y, k*Δ*t*), where Δ*x* = 1/*M*_*x*_, Δ*y* = 1/*M*_*y*_, Δ*t* = *T /M*_*t*_, while *i* = 0, …, *M*_*x*_, *j* = 0, …, *M*_*y*_, and *k* = 0, …, *M*_*t*_. We further simplify the notation for the spatial part as Ξ = {(*i*Δ*x, j*Δ*y*) |*i* = 0,…, *M*_*x*_, *j* = 0,…, *M*_*y*_}. We will use 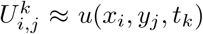 to denote the discretized approximation at (*x*_*i*_, *y*_*j*_, *t*_*k*_), and similarly, 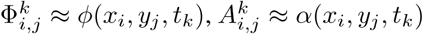.

For a sufficiently small Δ*t* > 0, a first-order approximation of *f* ((*k* + 1)Δ*t*; *a*), *N* ((*k* + 1)Δ*t*; *a*) starting from (*f* (*k*Δ*t*), *N* (*k*Δ*t*)) = (*x*_*i*_, *y*_*j*_) for *any k* ∈ {0, …, *M*_*t*_}, with a control value *a* ∈ {0, 1} is

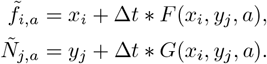

Let 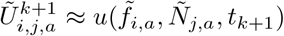. Since Eq. (S7.1.1) is in a Mayer form [7], the discretized dynamic programming equation is

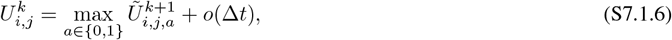

where 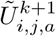 is evaluated by a bi-linear interpolation using the U values from the 4 neighboring gridpoints surrounding 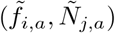. Note that 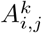 is found as the argmax of Eq. (S7.1.6) at each gridpoint. This straightforward time-marching scheme is summarized in Algorithm S1. In all of our numerical experiments, we have used *M*_*x*_ = *M*_*y*_ = 1600 on each side of the unit *fN* -square, and Δ*t* = 6.25 *×* 10^−3^. *To obtain the numerical solution for constitutive killers, one can simply apply Algorithm S1 with a* = 1 *without the maximization*.

#### Remark III

Despite the time-dependent nature of the problem, one is typically interested in the 0-th time slice of the value function (i.e., *u*(*x, y*, 0)), which predicts the value at *t* = *T* starting from *t* = 0 for all possible initial states. We note that by setting a sufficiently large *T* in Algorithm S1, any *k*-th time slice serves as the 0-th slice for a reduced horizon of *T* − *t*_*k*_. This allows us to obtain the prediction for a range of horizons simultaneously in a single sweep.

#### Algorithm S1

Finite-horizon value function computation

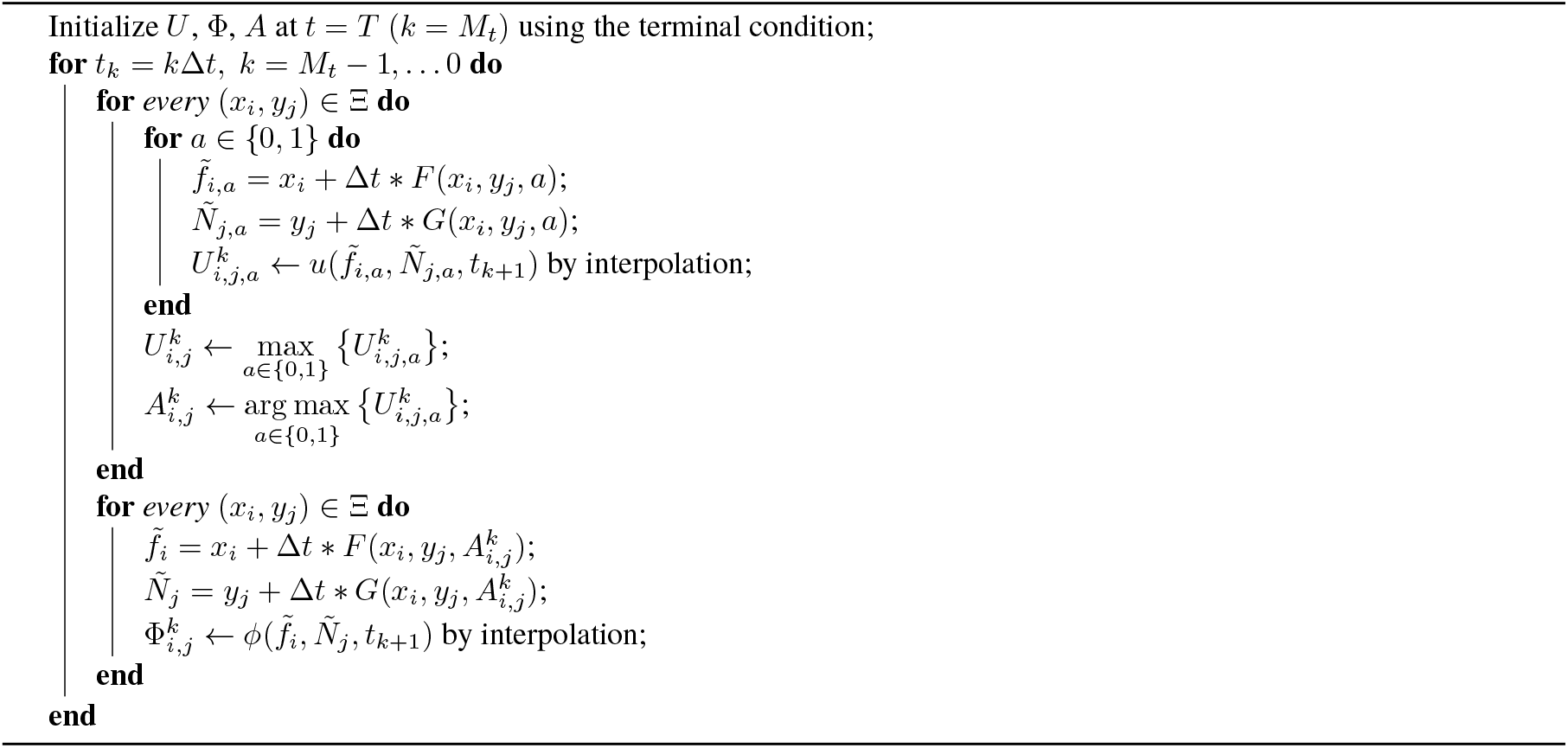

### S7.2 Approximating population limits under competitions

When sensitives and killers compete under regular dilutions, we are interested in whether they can coexist or if one population will dominate the other. As mentioned in the main text, this can be numerically found by a repetitive mapping using *u*(*x, y*, 0) and *ϕ*(*x, y*, 0) computed by Algorithm S1.

Let 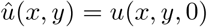 and 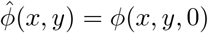. We will use 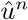 and 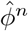 *s* to denote the relative abundance of the killer, and the normalized total population, respectively, by the end of *n*-th cycle. Thus, by definition, 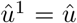 and 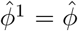. Based on our assumption, after the *n*-th dilution, *f* ((*nT*)^+^) = *f* ((*nT*)^−^) and *N* ((*nT*)^+^) = *ρN* ((*nT*)^−^). It follows that

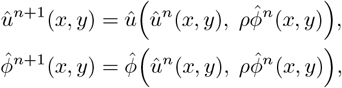

with *f* (0) = *x* and *N* (0) = *y* and following policy *α*. We repeat the process until 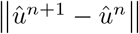 is small, and output 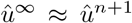 and 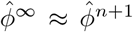 as the limits. Given our focus on this limit, we describe *α* as a “myopic” policy. This designation highlights that *α* is only optimal for a single period. The truly optimal policy for an infinite number of periods would typically adapt from one period to the next. Our full method of approximating 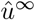 with this “myopic” policy is summarized in Algorithm S2 with the usual notations of solution on the discretized grid: 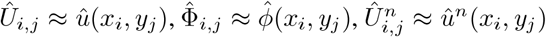 and 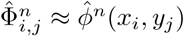.

#### Algorithm S2

Limiting performance of the myopic policy

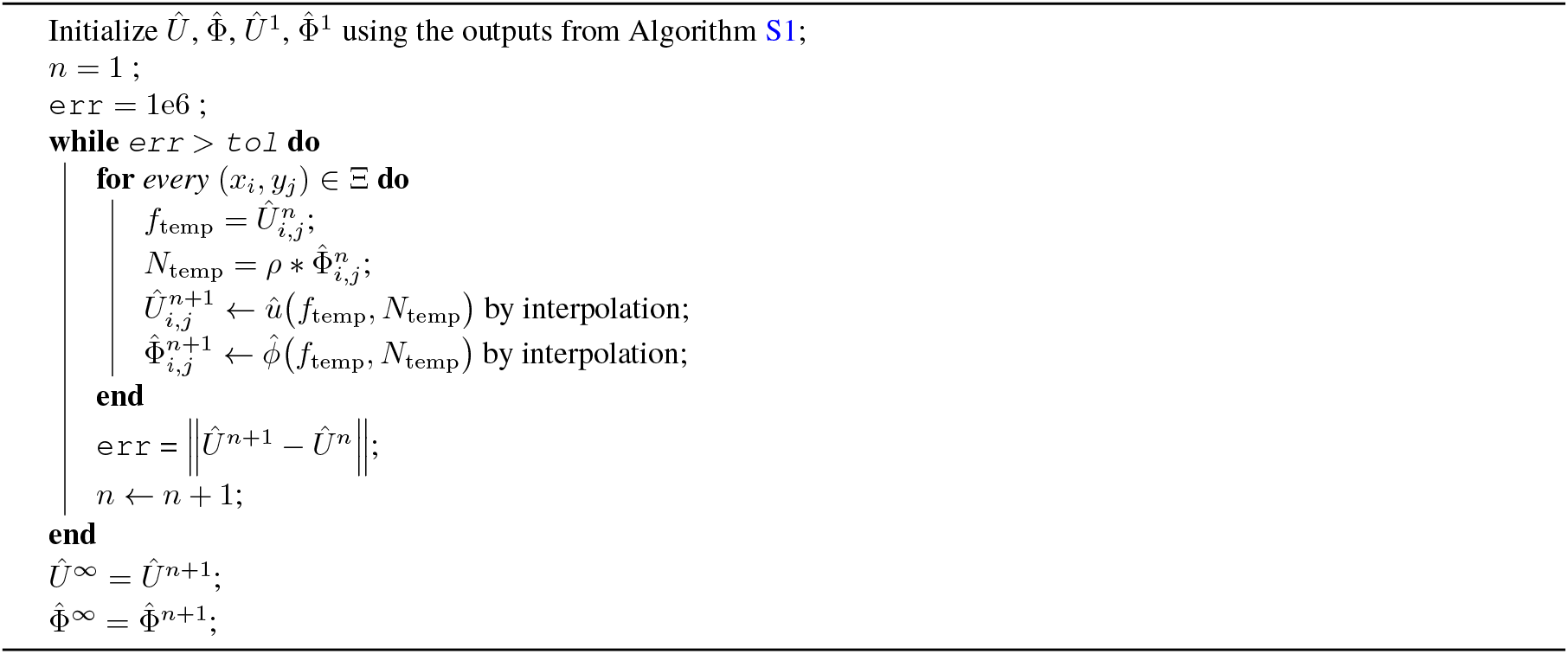

#### Remark IV

Given our interest in the limit as the number of dilutions approaches infinity, we can significantly accelerate Algorithm S2. Rather than updating just one cycle with 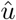 and 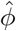 per iteration, we can exponentially increase the number of cycles updated at each iteration. Specifically, at the *n*-th iteration, we can update 2^*n*^ cycles by using the results from the previous iteration. Let 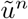 and 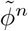 represent the values at the *n*-th iteration of this accelerated algorithm, starting with 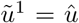 and 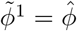, we now have

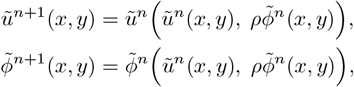

with 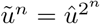 and 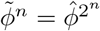. This accelerated algorithm is summarized in Algorithm S3.

#### Algorithm S3

Accelerated computation of 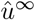

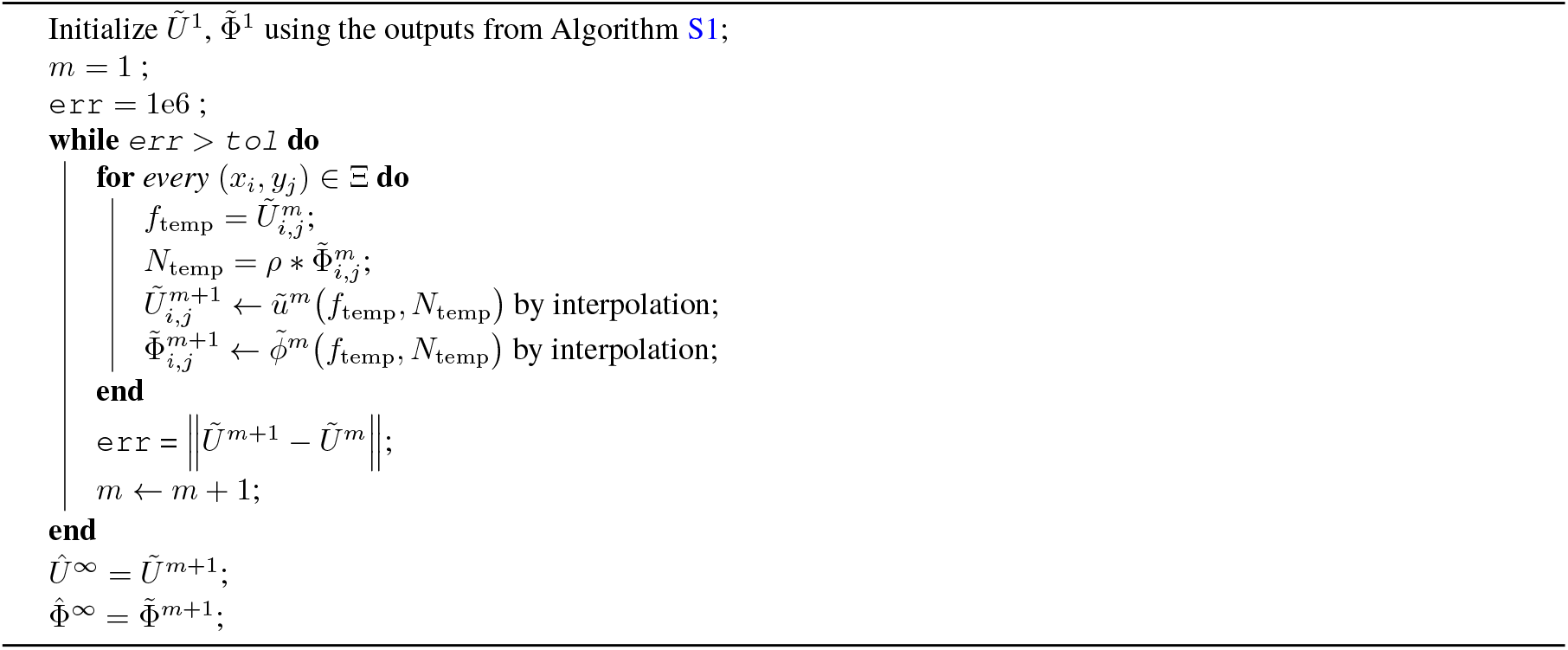

#### Remark V

Either Algorithm S2 or Algorithm S3 can be applied to each time-slice in Algorithm S1 to compute the 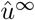 and 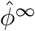 for a range of horizons simultaneously in a single sweep. This can be achieved by defining 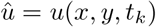 and 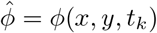 at each *k*-th slice, and thus obtaining the limits for each reduced horizon *T* − *t*_*k*_.

### S7.3 For the “strategically-optimal” killer

The HJB PDE (S6.1.1) associated with “tactically-optimal” killers can be numerically computed via standard Value [10–12] (or Value-Policy [13]) Iterations with a semi-Lagrangian discretization [5, 6, 9]. Here, we propose a similar Value-Policy Iterations (VPI) scheme to compute Eq. (S6.2.2) for the “strategically-optimal” killers.

Assuming the same spatial discretization Ξ as in §S7.1, we denote the approximate solution to the value function as *W*_*i,j*_ ≈ w(x_i_, y_j_). Similarly to §S7.1 (but now with *k* always fixed at 0), the foot of the characteristics starting from a gridpoint (*x*_*i*_, *y*_*j*_) reaches a new state 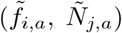 for a sufficiently small Δ*t* with control value *a*.

Therefore, from the Dynamic Programming Principle (DPP), we have

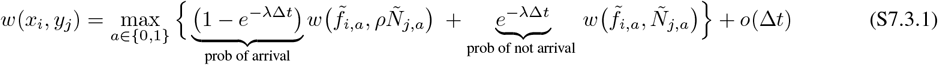

yielding the discretized version

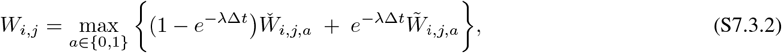

where 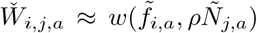 and 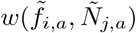 are computed through a *bi-linear* interpolation of the *W* values from the four neighboring gridpoints surrounding 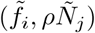 and 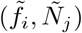, respectively. The optimal feedback policy A_*i,j*_ ≈ *α*(*x*_*i*_, *y*_*j*_) is recovered as an argmax in Eq. (S7.3.2).

We start with value iterations where we solve the nonlinear Eq. (S6.2.2) by a Gauss-Seidel relaxation. Let 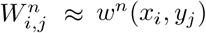 and 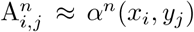 be the discretized solution/policy at the *n*-th iteration at gridpoint (*x*_*i*_, *y*_*j*_). We use err to denote the *L*_∞_-norm of *W* -change in the current value iteration. Whenever `err` stagnates, we proceed to the “*policy-evaluation*” (PE) step.

In the PE step, we compute the value function by solving a system of linear equations with a fixed policy 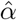 (recovered from the most recent value iteration). A first-order approximation of the system Eq. (2) starting from (*x*_*i*_, *y*_*j*_) with policy 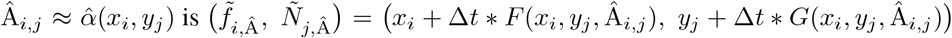. We thus solve a linear system of equations

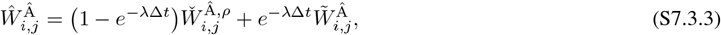

where 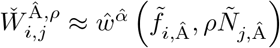 and 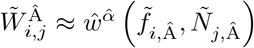 are again computed through a bi-linear interpolation.

After obtaining the solution to Eq. (S7.3.3), we return to the value iteration part and repeat the process until `err < tol`, where tol is a preset tolerance of convergence. In all of our numerical experiments, we have used *M*_*x*_ = *M*_*y*_ = 1600 on each side of the unit *fN* -square, Δ*t* = 0.025, and tol = 10^−6^. our full method is summarized in Algorithm S4.

Under mild technical assumptions, Kushner and Dupuis [14, Chapters 10 & 16] showed that the discretized solution derived from a general jump-diffusion process converges to the value function using standard iterative methods. Our model forms a PDMP, which is just a specific case of jump-diffusion processes.

#### Algorithm S4

Value-Policy Iterations for the non-local HJB equation Eq. (S6.2.2)

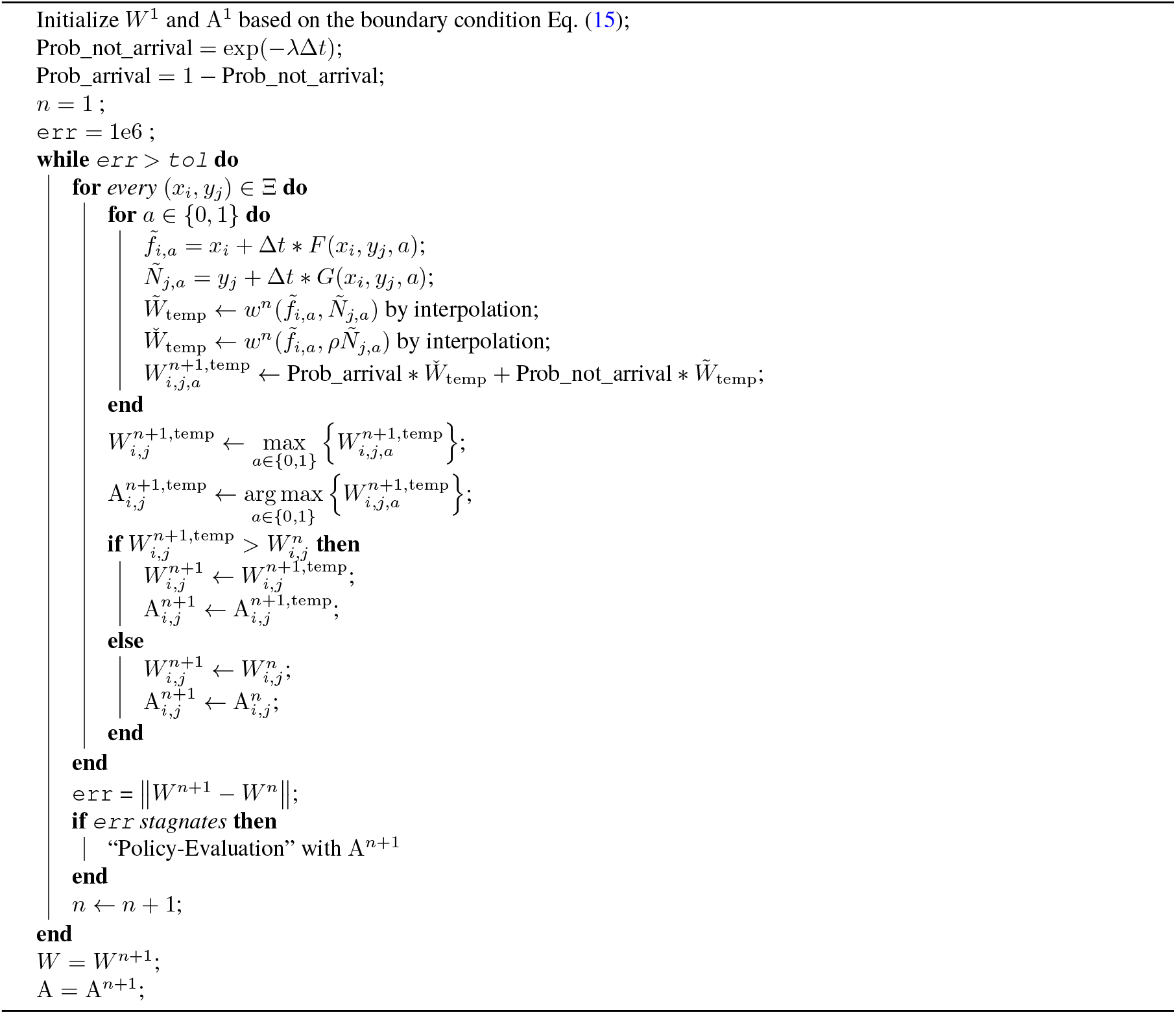

#### Remark VI

This is a semi-Lagrangian-based method for solving the linear non-local probabilistic performance metric Eq. (9) in the main text, we will apply it to assess the probability performance of *α*_0_ and *α*_1_.

## S8 Population-dependent (hyperbolic) win and defeat boundaries

In the main text, we have focused on *fraction-dependent (vertical)* boundary arising from the definitions of killers’ victory and defeat:

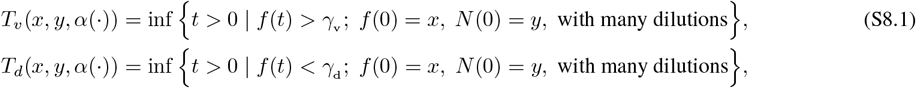

where both stopping criteria depended on the *fraction* of the killer strain, *f* (*t*), only. In the main text, we have used *γ*_v_ = 0.99 and *γ*_d_ = 0.01.

However, there are many other suitable ways of defining killers’ victory/defeat. In this section, we explore *population-dependent (hyperbolic)* win and defeat boundaries and demonstrate that the results remain qualitatively similar to those obtained in the main text.

Mathematically, we now define the (random) victory/defeat time as

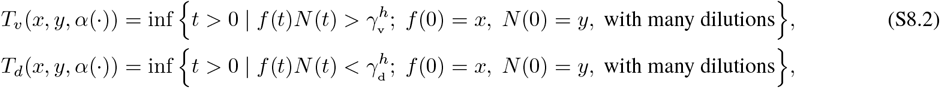

so that the stopping criteria depend on the *population size* of the killer strain, *n*_K_(*t*) = *f* (*t*)*N* (*t*).

As a result, while the equations for the “strategically-optimal” killer (*w*(*x, y*)) and the probabilistic performance metric 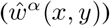 remain unchanged, their respective boundary conditions are now specified on two *hyperbolas*:

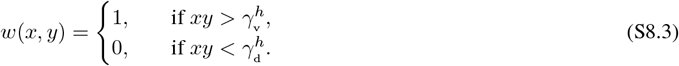

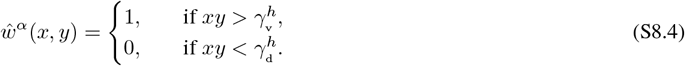

Let *w*_*v*_ denote the value function with vertical boundaries and *w*_*h*_ the one with hyperbolic boundaries. Using the same parameter values as in Fig. S7 in the main text but with 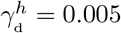 and 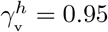, we find that *α*_∞_ computed with hyperbolic boundaries (Fig. S8B) is mostly the same as in Fig. S7B, except for the bottom right corner of the orange region. Consequently, the value function remains qualitatively the same too. To quantify the difference between *w*_*v*_ (Fig. S7A) and *w*_*h*_ (Fig. S8A), we calculate the mean absolute difference

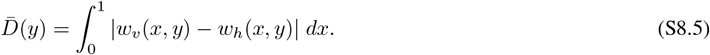

and plot it for all initial populations *y* ∈ [0, 1]. Fig. S8C shows that 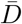is only large when *y <* 0.05 due to a more abrupt transition from zero to a positive winning probability in *w*_*v*_ compared to *w*_*h*_ when *x* is close to 1 and *y* is small. This is not surprising since these initial conditions are much closer to the hyperbolic defeat boundary 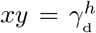 than they are to the vertical defeat boundary *x* = *γ*_d_. For *y* > 0.05, this *x*-averaged difference is very close to zero, suggesting that for most initial conditions the probability of killers’ winning is largely insensitive to the type of boundary used.

**Figure S8.**
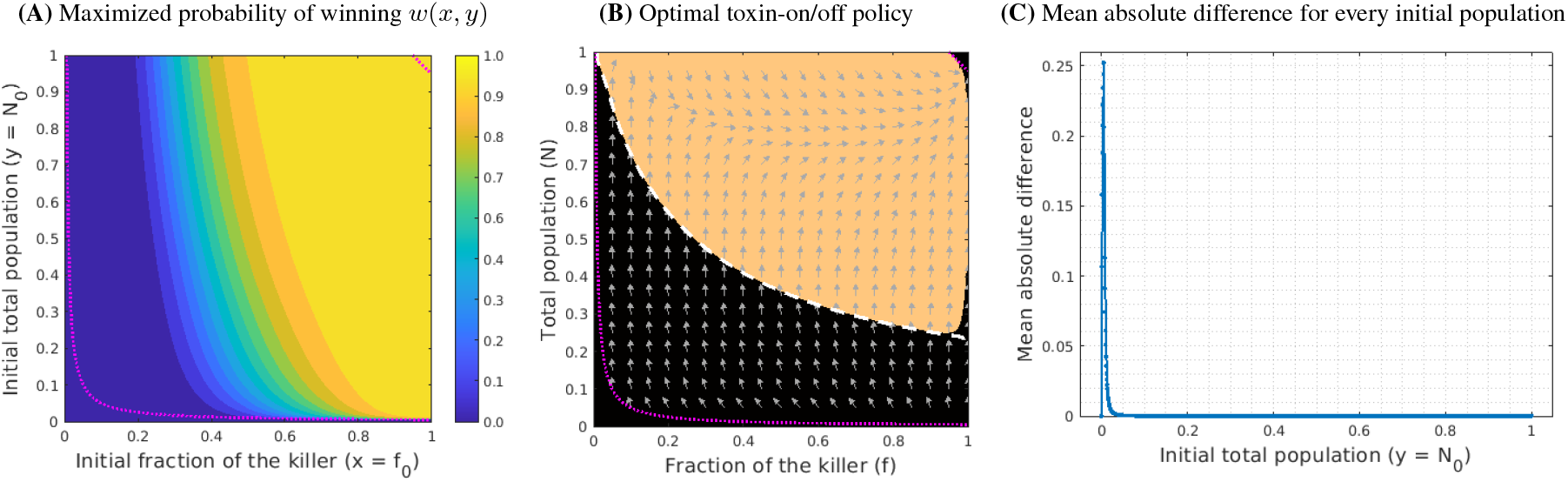
“Strategically-optimal” killers with hyperbolic win/defeat boundaries. The optimal toxin-on region (orange in panel B) is almost the same as the one computed with vertical boundaries in Fig. S7B. (The toxin-on/off switch curve from the latter is shown here as a white-dashed line). As a result, the maximized probability of winning (panel A) is also very similar to the one computed with vertical boundaries in Fig. S7A. Panel C shows the *x*-averaged mean absolute difference between panel A and Fig. S7A across all initial populations *N*_0_ = *y* ∈ [0, 1]. This difference is only noticeable when *y <* 0.05. In (B), gray arrows denote the vector field directions corresponding to Eq. (2) in the main text with *a* = *α*_∞_ (*f, N*) computed using hyperbolic win/defeat boundaries. In all panels, the victory and defeat barriers (*γ*_v_ and *γ*_d_, respectively) are plotted with a magenta dotted line. All parameter values are the same as in Fig. S7.

This conclusion also generally holds true when using the above hyperbolic boundaries to compute 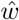 with *α*_0_ and *α*_1_. Analogously to Fig. 8 in the main text, we again focus on the initial condition (*f*_0_, *N*_0_) = (0.5, 0.1) and compare the performance of these toxin-production policies for a range of (*ρ, λ*) values. Fig. S9 shows that these heat maps are both qualitatively and quantitatively similar to those in Fig. 8.

We similarly quantify the boundary-related difference in the probabilistic performance of constitutive and tactically-optimal killers in Fig. S10. Let 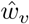 and 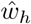 denote the probabilistic performance with vertical and hyperbolic victory/defeat boundaries, respectively. Focusing on the same initial condition (*f*_0_, *N*_0_) = (0.5, 0.1), we observe that 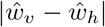 is negligible across half of the heat map (*ρ* ≥ 0.6) for both *α*_∞_ and *α*_1_. In these two cases, a noticeable difference (with a maximum of approximately 0.017) is observed when the dilutions are strong (*ρ* ≤ 0.55). For *α*_0_, the maximum difference is slightly lower, around 0.014, while the region with noticeable differences is larger (*ρ* ≤ 0.6). This is expected, as a stronger dilution more likely leads to a defeat under hyperbolic boundaries (due to a significantly larger Δ_d_) compared to vertical boundaries. These results indicate that our observations and conclusions in the main text remain largely unaffected by whether the victory or defeat of the killer is defined by its fraction or population.

**Figure S9.**
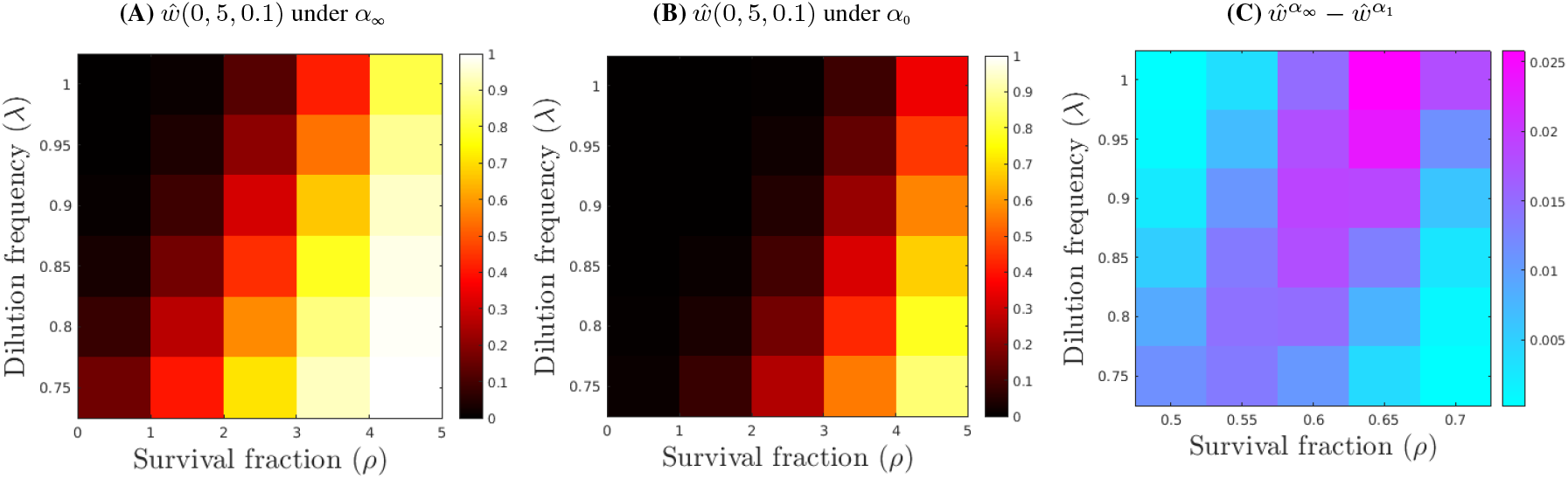
Hyperbolic boundaries: comparison of probabilistic performance for different types toxin-production policies starting from (*f*_0_, *N*_0_) = (0.5, 0.1) for a range of dilution strengths and frequencies. Policy *α*_1_ is recomputed for each *λ*, while policy *α*_∞_ is recomputed for each (*ρ, λ*) combination. The results remain both qualitatively and quantitatively similar to Fig. 8 in the main text. A stronger survival rate (larger *ρ*) combined with less frequent dilutions (smaller *λ*) increases the chances of toxin-producers winning for all three policies. It is clear that the strategically optimal (panel A) and tactically optimal killers significantly outperform the constitutives (panel B). The differences in 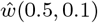 between *α*_∞_ and *α*_1_ are still small, with the discrepancy increasing toward the upper right corner (panel C).

**Figure S10.**
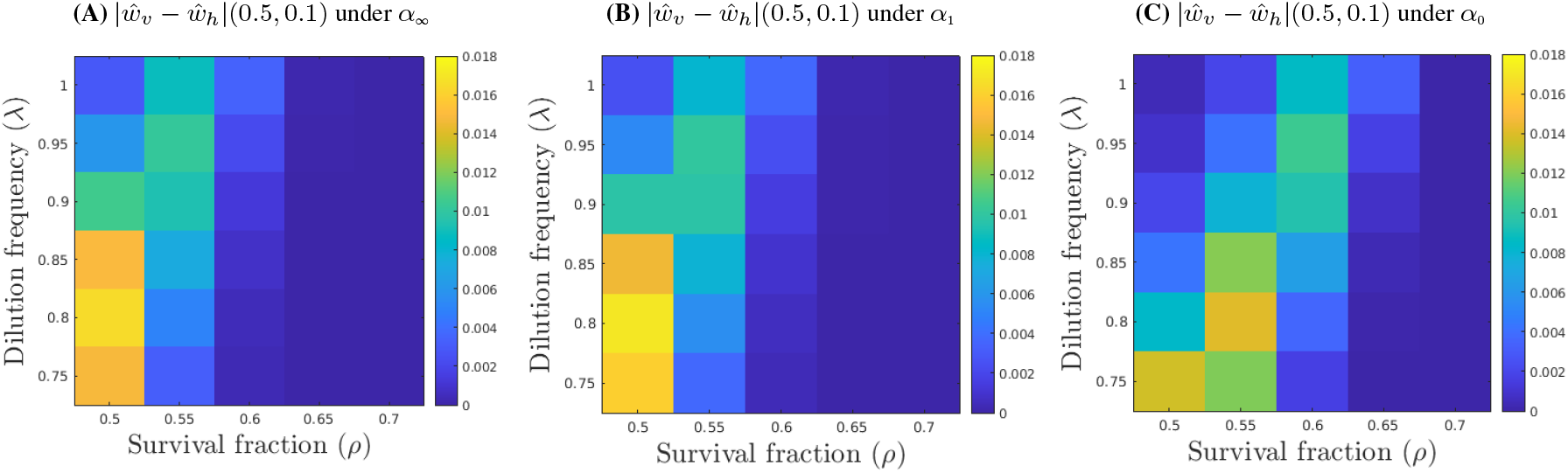
Hyperbolic boundaries: absolute difference in 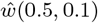 resulting from two types of boundaries computed for a range of *ρ* and *λ* values. The differences under *α*_∞_ (panel A) and *α*_1_ (panel B) are again similar, with a maximum difference of around 0.017 when *ρ* = 0.5. For *α*_0_ in panel C, the region of noticeable differences is slightly larger (*ρ* ≤ 0.6) although the maximum difference remains relatively small (≈ 0.014). All three panels share the same colorbar.

## S9 Monte Carlo simulations with “Binomial dilutions”

The results in the main text are all produced under “deterministic” dilution outcomes. That is, after each dilution, the relative abundances are preserved while only a *ρ* proportion of the total population survives. In this section, we present results under a specific form of *random* dilution outcomes and demonstrate, using Monte Carlo (MC) simulations, that they are qualitatively similar to the previous results.

In particular, we adopt a “Binomial Sampling” strategy to produce random dilution outcomes, which we refer to as “Binomial dilutions.” Let *f* ^−^ be the pre-dilution fraction of the killers, and *N* ^−^ be the pre-dilution total population. Accordingly, the actual pre-dilution number of killer cells is 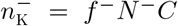, and the pre-dilution number of sensitive cells is 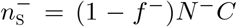. We assume each cell has an independent survival probability *ρ* after each dilution. As a result, the post-dilution number of cells is a *Binomial random variable*:

- Post-dilution number of killer cells: 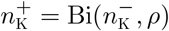;
- Post-dilution number of sensitive cells: 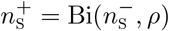.

Consequently, the random post-dilution (normalized) total population is 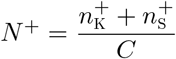, and the random post-dilution fraction of killers is 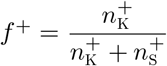, which will serve as the initial condition for the next cycle.

We first conduct Monte Carlo simulations on a uniform grid. Starting from each (*x*_*i*_, *y*_*j*_) = (*i*/10, *j*/10) with *i, j* = 1, 2,… 9, each sample was simulated with *n* = 200 dilutions. Fig. S12 shows the empirical distributions of *f* ((*nT*)^−^) with *T* = 1 and Fig. S11A shows their respective means. We observe that most distributions are unimodal, being either nearly 0 (all sensitives) or nearly 1 (all killers). The two exceptions with a bimodal distribution, peaking at 0 or 1, intersect precisely with the boundary (black-dashed line in Fig. S11A) that separates the initial conditions leading to a *deterministic* victory of the killers under proportional dilutions. Additionally, the killer-winning region (dark-red background in Fig. S11A) under “Binomial dilutions” aligns well with this “deterministically-killer-winning” region shown in Fig. 4C in the main text. This is not surprising since the stochastic fluctuations introduced by “Binomial dilutions” can be sufficiently large to cause samples starting near the boundary to drift towards either competitive exclusion over successive dilutions. See Fig. S11C for such an example starting from (*f*_0_, *N*_0_) = (*x, y*) = (0.5, 0.4). However, these fluctuations are typically insufficient to alter the fate of samples starting further from the boundary, where initial conditions strongly favor one strain.

Considering that focusing on a single initial condition for all samples might not capture enough information, we conducted additional Monte Carlo simulations using “Binomial dilutions” with *uniformly random in a cell* initial conditions. Specifically, for each grid cell centered at (*x*_*i*_, *y*_*j*_), the initial condition for each sample was chosen uniformly at random from the square (*f*_0_, *N*_0_) = (*x, y*) ∈ [*x*_*i*_ − 0.05, *x*_*i*_ + 0.05] *×* [*y*_*j*_ − 0.05, *y*_*j*_ + 0.05]. This strategy diversifies the range of initial conditions, increasing the likelihood of intersecting the boundary of the “deterministically-killer-winning” region. As a result, we see from Fig. S11B that almost all cells intersecting or near the dashed-line boundary now exhibit intermediate mean values. Moreover, Fig. S13 shows that these cells again have a bimodal distribution, with random dilution outcomes pushing the dynamics towards one of the two competitive exclusions. Comparing Fig. S13 with Fig. S12, we further observe that the median at each grid cell remains unchanged. This consistency underscores the robustness of our simulation results, indicating that the conclusions in the main text would largely remain valid even under “Binomial dilutions.”

**Figure S11.**
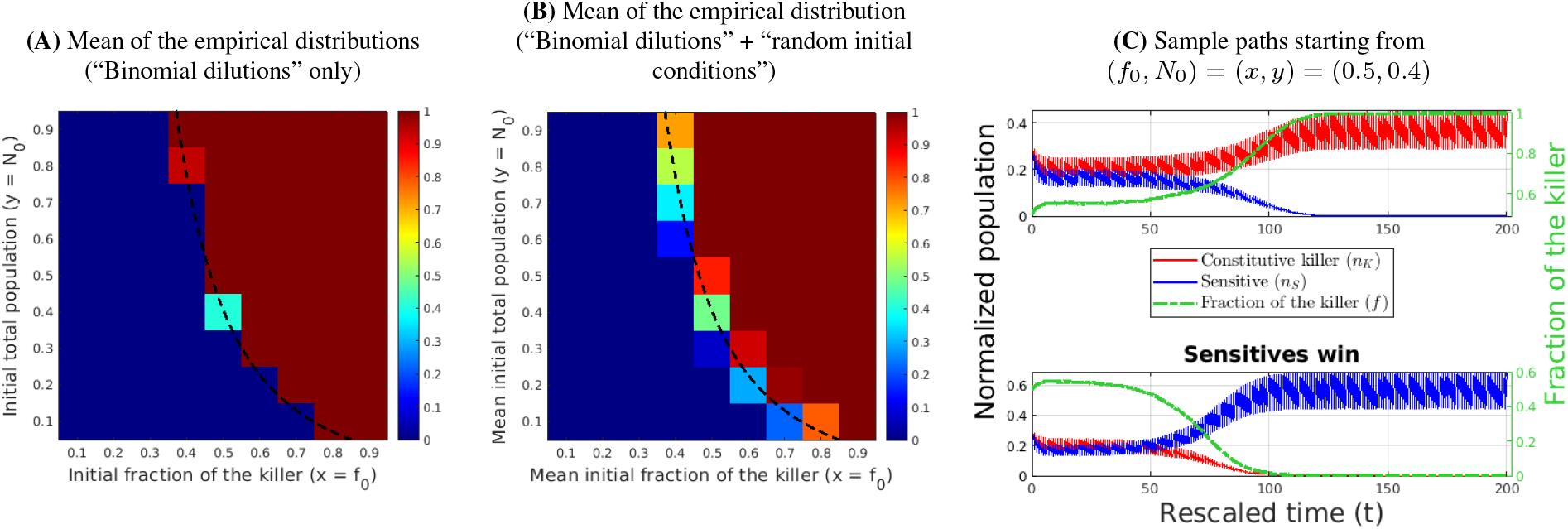
Monte Carlo simulations with “Binomial dilutions” on a uniform cell grid. (A) The mean of the empirical distribution, sampled with “Binomial dilutions” starting from the center of each cell in (*f*_0_, *N*_0_) = (*x, y*) space. The resulting distribution is almost always unimodal (dark blue - all sensitives; dark red – all killers). The exceptions are seen in only two cells among those intersected by the boundary (shown by a black dashed line) that separates the initial conditions leading to a deterministic victory of the killers under proportional dilutions (cf. Fig. 4C in the main text). (B) Most means of the empirical distribution, sampled with both “Binomial dilutions” and “uniformly random in a cell” initial conditions, are also close to 0 or close to 1 in most cells. However, most cells that intersect or are close to that dashed line boundary now have more diverse intermediate mean values. In both cases, such cells exhibit a bimodal distribution with peaks at 0 and 1; see Figs. S12&S13 for the actual distributions. Panel (C) shows two sample trajectories starting from (*f*_0_, *N*_0_) = (*x, y*) = (0.5, 0.4) resulting in different competitive exclusion outcomes due to the randomness in Binomial dilutions. All Monte Carlo simulations were conducted with 10^5^ samples and 200 dilutions using parameter values *T* = 1, *ε* = 0.2, *r*_KS_ = 0.85, *γ* = 1, and *ρ* = 0.65. In (A), all samples for each grid cell start from the same initial condition (*x*_*i*_, *y*_*j*_) = (*i*/10,*j*/10) with *i, j* = 1,…, 9. In (B), for each grid cell centered at (*x*_*i*_, *y*_*j*_), the initial condition for each sample was chosen uniformly at random from the square (*f*_0_, *N*_0_) = (*x, y*) ∈ [*x*_*i*_ − 0.05, *x*_*i*_ + 0.05] *×* [*y*_*j*_ − 0.05, *y*_*j*_ + 0.05].

**Figure S12.**
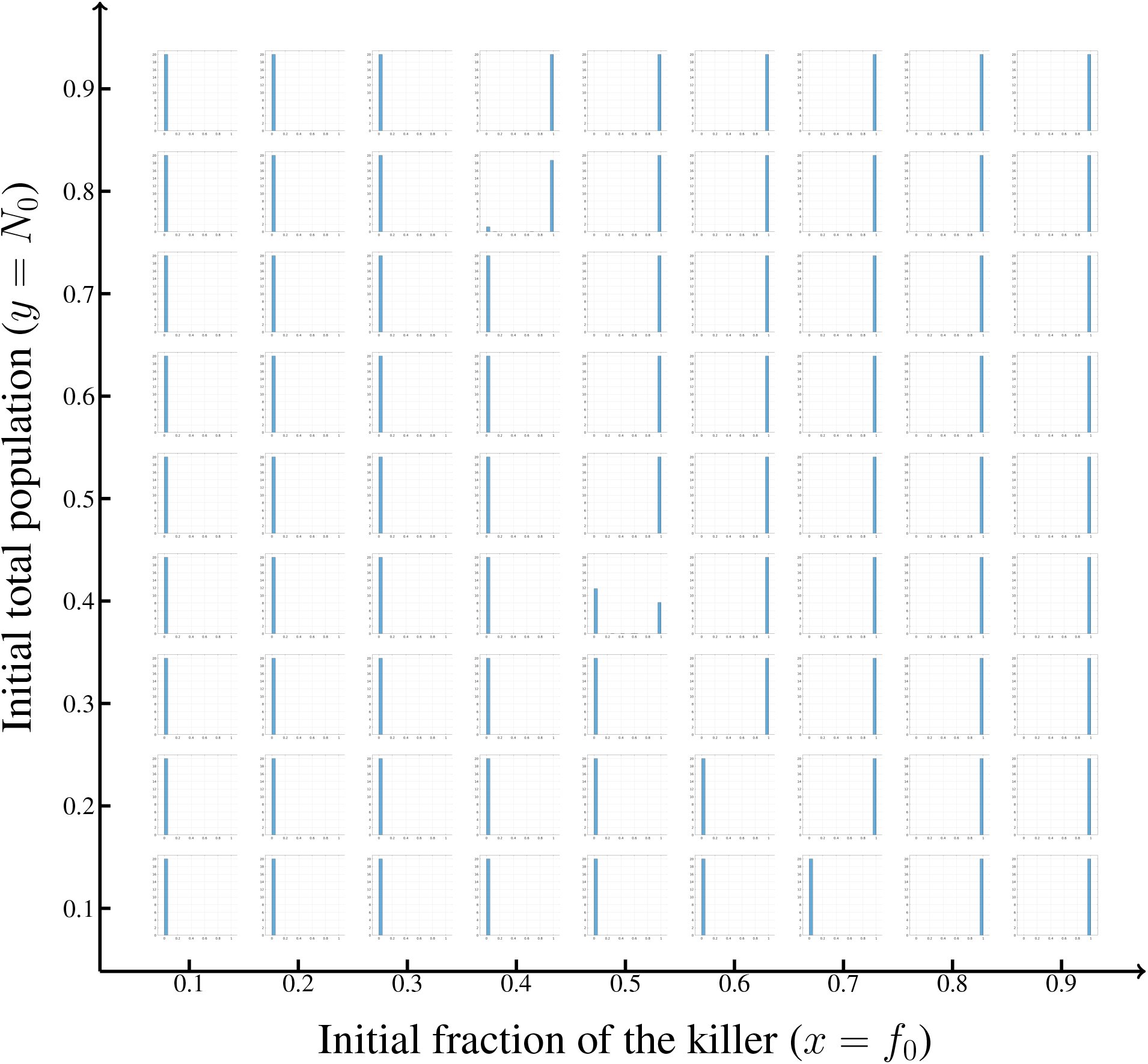
Empirical distributions of the fraction of killers with “Binomial dilutions” on a uniform grid. Most of the distributions are unimodal (either almost entirely 0 or almost entirely 1), except for two that are bimodal. The horizontal axis (of the entire figure) represents the initial fraction of the killer while the vertical axis encodes the initial total population for the simulations. For each panel, an empirical distribution of *f* (starting from the same (*x*_*i*_, *y*_*j*_) = (*i*/10,*j*/10) with *i, j* = 1,…, 9) after 200 dilutions is shown by a histogram. All panels share the same horizontal and vertical axes. The parameter values are the same as in Fig. S11A.

**Figure S13.**
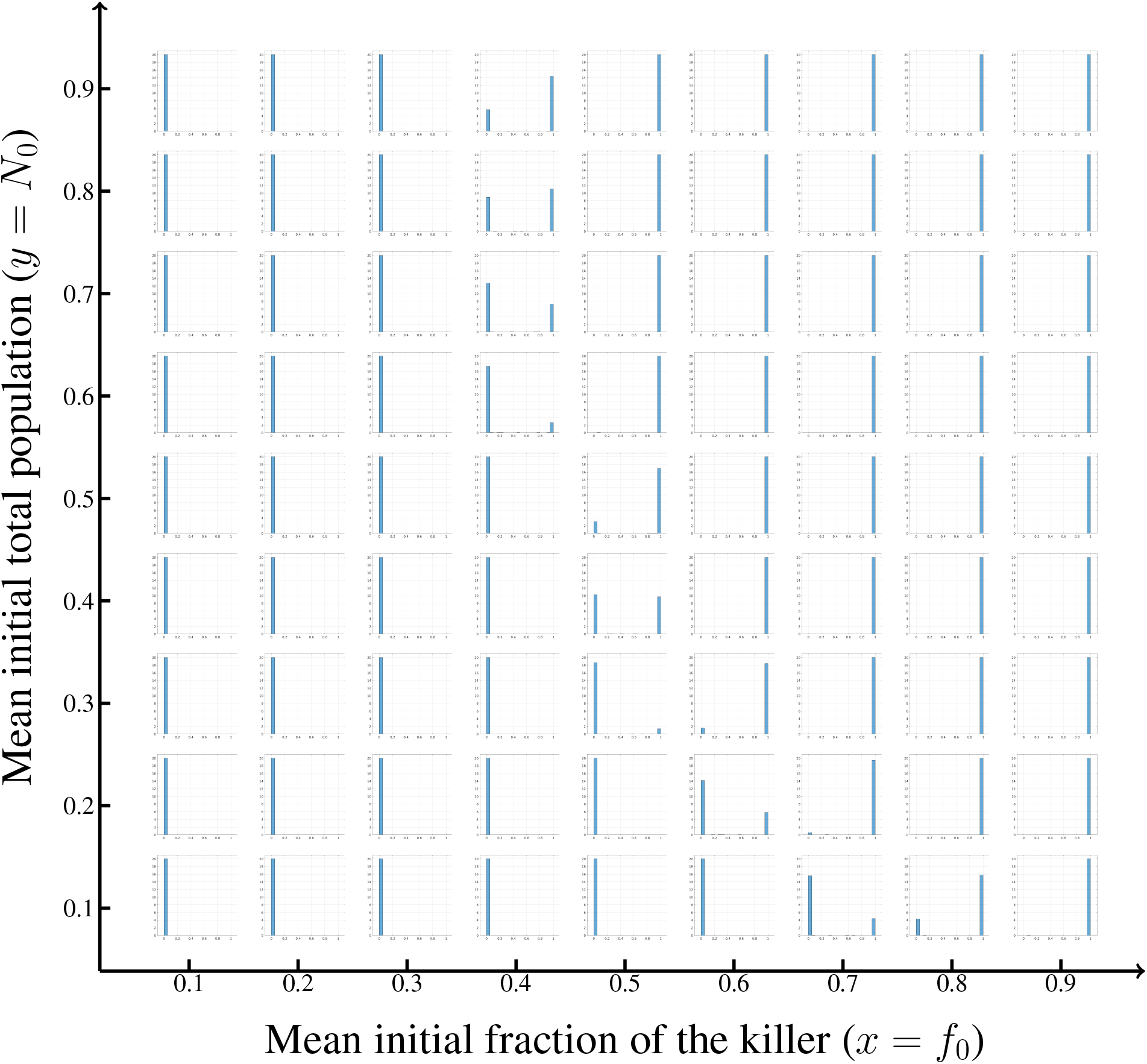
Empirical distributions of the fraction of killers with “Binomial dilutions” and “uniformly random in a cell” initial conditions on a cell grid. Most of the distributions are unimodal (either almost entirely 0 or almost entirely 1). However, the ones near the boundary of the “deterministically-winning” region (depicted as a black-dashed line) are *bimodal*. The horizontal axis (of the entire figure) represents the mean initial fraction of the killer while the vertical axis encodes the mean initial total population for the simulations. For each panel, an empirical distribution of *f* (*t*) after 200 dilutions, with the initial condition chosen uniformly at random within the grid cell centered at (*x*_*i*_, *y*_*j*_) = (*i*/10,*j*/10) with *i, j* = 1,…, 9, is shown by a histogram. All panels share the same horizontal and vertical axes. The parameter values are the same as in Fig. S11B.

The results presented above appear to be qualitatively similar for all tested parameter values. As the penalty for the ability to produce toxin and the penalty for actually producing it decreases (i.e., *r*_KS_ → 1 and *ε* → 0), it becomes more difficult for the sensitives to win and the dark blue region in Fig. S11A significantly shrinks; e.g., Fig. S14A. Nevertheless, when starting in the overwhelming majority, the sensitives can occasionally win under “Binomial dilutions” even if *r*_KS_ = 1; see Fig. S14A. I.e., Binomial dilutions might wipe out even super-competitive killers if they are added in small numbers to an established population of sensitives. If the Monte-Carlo simulations are started from randomly chosen initial conditions, the part of the (*f*_0_, *N*_0_) = (*x, y*) domain where both strains have a chance to win increases; see Fig. S14B. This observation aligns well with the results obtained for *r*_KS_ < 1 (Fig. S11B), confirming the strong dependence of competitive exclusion on initial conditions, even when *r*_KS_ = 1.

**Figure S14.**
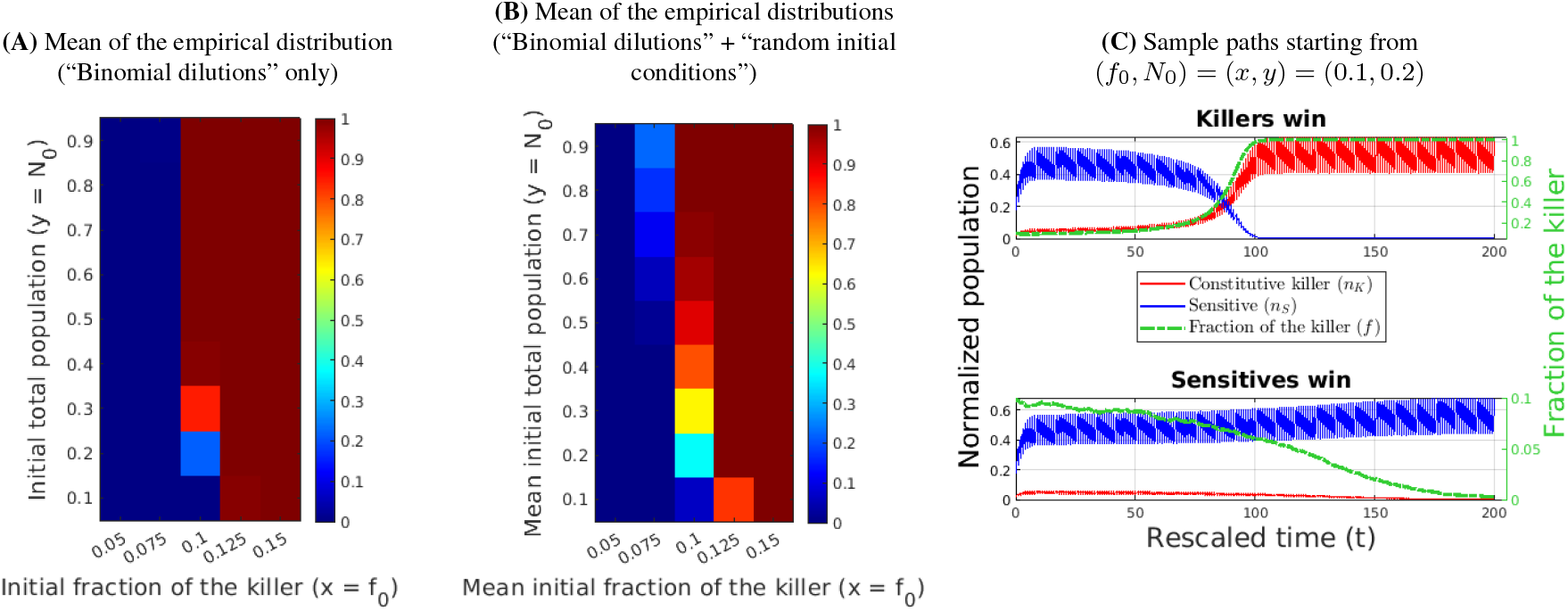
Monte Carlo simulations with “Binomial dilutions” on a uniform cell grid with *r*_KS_ = 1 and *ε* = 0.1. (A) The mean of the empirical distribution, sampled with “Binomial dilutions” starting from the center of each cell in (*f*_0_, *N*_0_) = (*x, y*) space. The killers will always win (dark red) if *f*_0_ *>* 0.1 while the sensitives will always win (dark blue) if *f*_0_ < 0.1. Bimodal distributions, peaking at 0 or 1, occur for *f*_0_ = 0.1. (B) When sampling with both “Binomial dilutions” and “uniformly random in a cell” initial conditions, more bimodal distributions with more diverse intermediate mean values appear near *f*_0_ = 0.1. Panel (C) shows two sample trajectories starting from (*f*_0_, *N*_0_) = (*x, y*) = (0.1, 0.2) resulting in different competitive exclusion outcomes due to the randomness in Binomial dilutions. All Monte Carlo simulations were conducted with 10^5^ samples and 200 dilutions using parameter values *T* = 1, *ε* = 0.1, *r*_KS_ = 1, *γ* = 1, and *ρ* = 0.65. In (A), all samples for each grid cell start from the same initial condition (*x*_*i*_, *y*_*j*_) = (*i*/40,*j*/10), with *i* = 2,…, 6, and *j* = 1, 2,…, 9. In (B), for each grid cell centered at (*x*_*i*_, *y*_*j*_), the initial condition for each sample was chosen uniformly at random from the square (*f*_0_, *N*_0_) = (*x, y*) ∈ [*x*_*i*_ − 0.0125, *x*_*i*_ + 0.0125] *×* [*y*_*j*_ − 0.05, *y*_*j*_ + 0.05].

## Footnotes

1 Unlike in the experiments reported in Fig. 1 (where nutrients were added at the start and never replenished) and in Fig. 2 (where nutrients were replenished at each dilution), the fixed carrying capacity used in Eq. (1) reflects a more realistic assumption of continuous nutrient supply.

2 We focus here on such “strictly proportional” dilutions for the sake of simplicity and computational efficiency. The results in SI Appendix §S9 show that for most initial conditions the conclusions remain largely the same even with a probabilistic dilution model, where each cell has probability *ρ* of surviving each dilution.

3 Our use of dynamic programming ensures that the computed policy is *globally* optimal and is obtained “in feedback form”. These are important advantages over the *Pontryagin Maximum Principle* approach, which is more commonly used in biological applications [66].

4 It is similarly possible to declare the victory/defeat criteria in terms of strain populations rather than fractions. For most initial conditions, this does not affect the qualitative policy and victory probabilities; see SI Appendix S8.

